# Oligopaint DNA FISH as a tool for investigating meiotic chromosome dynamics in the silkworm, *Bombyx mori*

**DOI:** 10.1101/2021.04.16.440181

**Authors:** Leah F. Rosin, Jose Gil, Ines A. Drinnenberg, Elissa P. Lei

## Abstract

Accurate chromosome segregation during meiosis is essential for reproductive success. Yet, many fundamental aspects of meiosis remain unclear, including the mechanisms regulating homolog pairing across species. This gap is partially due to our inability to visualize individual chromosomes during meiosis. Here, we employ Oligopaint FISH to investigate homolog pairing and compaction of meiotic chromosomes in a classical model system, the silkworm *Bombyx mori*. Our Oligopaint design combines multiplexed barcoding with secondary oligo labeling for high flexibility and low cost. These studies illustrate that Oligopaints are highly specific in whole-mount gonads and on meiotic chromosome spreads. We show that meiotic pairing is robust in both males and female meiosis. Additionally, we show that meiotic bivalent formation in *B. mori* males is highly similar to bivalent formation in *C. elegans*, with both of these pathways ultimately resulting in the pairing of chromosome ends with non-paired ends facing the spindle pole and microtubule recruitment independent of the centromere-specifying factor CENP-A.

**Author’s Summary:** Meiosis is the specialized cell division occurring exclusively in ovaries and testes to produce egg and sperm cells, respectively. The accurate distribution of chromosomes (the genetic material) during this process is essential to prevent infertility/sterility and developmental disorders in offspring. As researchers are specifically unable to study the mechanisms regulating meiosis in depth in humans, identifying broadly conserved aspects of meiotic chromosome segregation is essential for making accurate inferences about human biology. Here, we use a sophisticated chromosome painting approach called Oligopaints to visualize and study chromosomes during meiosis in the silkworm, *Bombyx mori*. We illustrate that Oligopaints are highly specific in *B. mori* and demonstrate how Oligopaints can be used to study the dynamics of meiotic chromosomes in diverse species.

## Introduction

Precise homolog pairing and unpairing during meiosis is essential for genetic recombination and accurate chromosome segregation. Errors in chromosome segregation during meiosis can lead to reduced fertility, miscarriages, or chromosomal disorders in progeny, such as Down Syndrome or Turner Syndrome (1). Decades of research has gone into characterizing the synaptonemal complex (SC), a proteinaceous structure that holds homologs together during meiotic prophase and is conserved across species (2, 3). Yet how homologs find each other and come together in 3D space is still poorly understood. One of the main reasons that homolog pairing has remained such an enigma is the lack of cytological tools available for assaying chromosome- and locus-specific pairing dynamics during meiosis. Several recent studies have taken advantage of advances in super resolution microscopy techniques, such as Structure Illumination Microscopy (SIM) and Stochastic Optical Reconstruction Microscopy (STORM), to visualize meiotic pairing in more detail than ever before (4–9). However, these approaches have been limited to studying pairing genome-wide by fluorescently labeling elements of the SC (5,7–12) or to visualizing small genomic loci by FISH (13–16).

Recent technological innovations in the design and synthesis of specialized DNA FISH probes called Oligopaints have made visualizing whole, individual chromosomes or complex sub- chromosomal loci in meiotic cells feasible. Unlike traditional BAC-based FISH probes, Oligopaints are computationally designed based on genome sequence data (17, 18). This approach allows for only unique, single copy sequences to be labeled, significantly increasing the specificity and resolution of FISH. Here, we leverage the flexibility of the Oligopaint design to add barcodes to label either whole chromosomes or different sub-chromosomal loci using the same set of oligos, as previously described (19). This multiplexed approach allows for many different highly specific FISH probes to be generated at low cost and high throughput. Oligopaints and related oligo-based FISH approaches have previously been used for karyotype analyses or characterization of interphase chromosome dynamics in *Drosophila*, *C. elegans*, mammals, and plants (16, 19–31). Recently, similar approaches have also been applied to the study of small chromosomal loci during meiosis (32, 33), but Oligopaints have never before been used to characterize compaction and pairing of multiple, whole chromosomes during meiosis. Finally, Oligopaints have never been used to visualize chromosomes in Lepidoptera (moths and butterflies).

Here, we combine Oligopaint DNA FISH with one of the first model systems ever used to study meiotic chromosomes, the silkworm moth *Bombyx mori*. *B. mori* are holocentric insects, with centromeres forming all along the chromosome during mitosis (34–37). The holocentric mitotic configuration is also seen in many plants and nematodes, including *C. elegans* (38–40). However, the holocentric chromosome configuration prevents accurate biorientation of bivalents formed after recombination and is therefore incompatible with canonical meiosis (40, 41). Instead, chromosomes in holocentric organisms often display “telokinetic” or “telokinetic-like” chromosomes during meiosis, where kinetochore activity is restricted toward telomere domains (42–48). In *C. elegans*, which telomere faces poleward to connect to the spindle microtubules is dictated by crossover position (42,46,49–51). A similar telokinetic mechanism for segregation meiotic chromosomes was also previously hypothesized to occur in *B. mori* (52–54) but has never before been directly observed. Furthermore, meiotic segregation in *C. elegans* occurs in the absence of the centromere-specifying factor Centromere Protein A (CENP-A) (51), and instead, microtubules either run parallel to chromosomes to facilitate segregation or directly penetrate chromosome ends (47, 55). Interestingly, CENP-A is entirely absent from the genomes of butterflies and moths (56). Yet, how moths and butterflies segregate chromosomes during meiosis in the absence of CENP-A remains to be explored.

Unlike *B. mori* spermatogenesis, which has been reported to support crossovers and canonical pairing, oogenesis in *B. mori* is quite unconventional. Chiasmata are not observed in female meiosis in silkworms and furthermore, the central elements of the SC break down just after pachytene (one of the sub-stages of meiotic prophase I) and the lateral elements of the SC are thought to be completely remodeled to form masses of “elimination chromatin” between the two homologs (54,57–59). This “elimination chromatin” or “modified SC” is reported to be over one micron in width, thereby ultimately undoing end-to-end homolog pairing while still holding homologs together until anaphase I (59). Thus, pairing along the entire length of the chromosomes is not expected after pachytene.

Our studies here illustrate that Oligopaints are robust and specific in germline cells and that Oligopaints can be used to visualize chromosomes even in unconventional model systems with draft genomes. Our FISH-based assays clearly demonstrate that telomeric regions face poleward and likely act as localized kinetochores during *B. mori* male meiosis and that both telomeres on any given chromosome harbor the ability to act as local kinetochores. Additionally, our data suggest that in female meiosis, homologs remain tightly paired throughout meiotic prophase I despite modifications to the SC. Overall, we provide the first extensive characterization of whole and sub-chromosome dynamics in meiosis in any species, thereby pioneering the use of Oligopaints as a tool for studying meiotic pairing and progression.

## Results

### *Bombyx mori* Oligopaint design

To visualize chromosomes in the silkworm, *B. mori*, we designed and generated Oligopaint libraries targeting six of the 27 autosomes and the Z sex chromosome. These Oligopaint libraries were designed using the Oligominer pipeline (18, 60) based on the updated 2019 silkworm genome assembly (61). Oligos were designed with 80 base pairs of homology and map to the genome only once (therefore only labeling unique, single copy sequences). This yielded a maximum probe density of approximately 3 oligos per kilobase (kb) of DNA. For most chromosomes, this density was then reduced to 1 or 1.5 probes per kb (see Table 1), a probe density that has been previously shown to be sufficient for whole chromosome paints (Rosin et al 2018). The resulting oligos are fairly evenly distributed along each chromosome, with gaps in regions where repetitive sequences are more abundant (Figure 1A-C). These oligo libraries were then multiplexed as previously described (19) with one or more barcode sequences to allow for the amplification of individual chromosomes, sub-chromosomal stripes, and/or active and inactive chromatin domains (Tables 1-5; Figure 1B-C, Figure S1). In total, the libraries consisted of 191,536 oligos (designated as “primary oligos”) up to 160 bp in length, which includes the 80 bp of homology, up to two unique 20 bp sub-chromosomal barcodes, and two 20 bp whole chromosome universal barcodes (Figure 1D). During the PCR amplification steps, secondary oligo binding sites are added to the primary oligos, to which fluorescently labeled secondary oligos will anneal during the FISH protocol (Figure 1E; (21, 30)). This method allows for increased flexibility when combining probes for multi-channel imaging. Together, this multiplexed probe design combined with secondary oligo labeling both increases the efficiency of Oligopaint synthesis and reduces the cost of Oligopaints.

**Figure 1.**
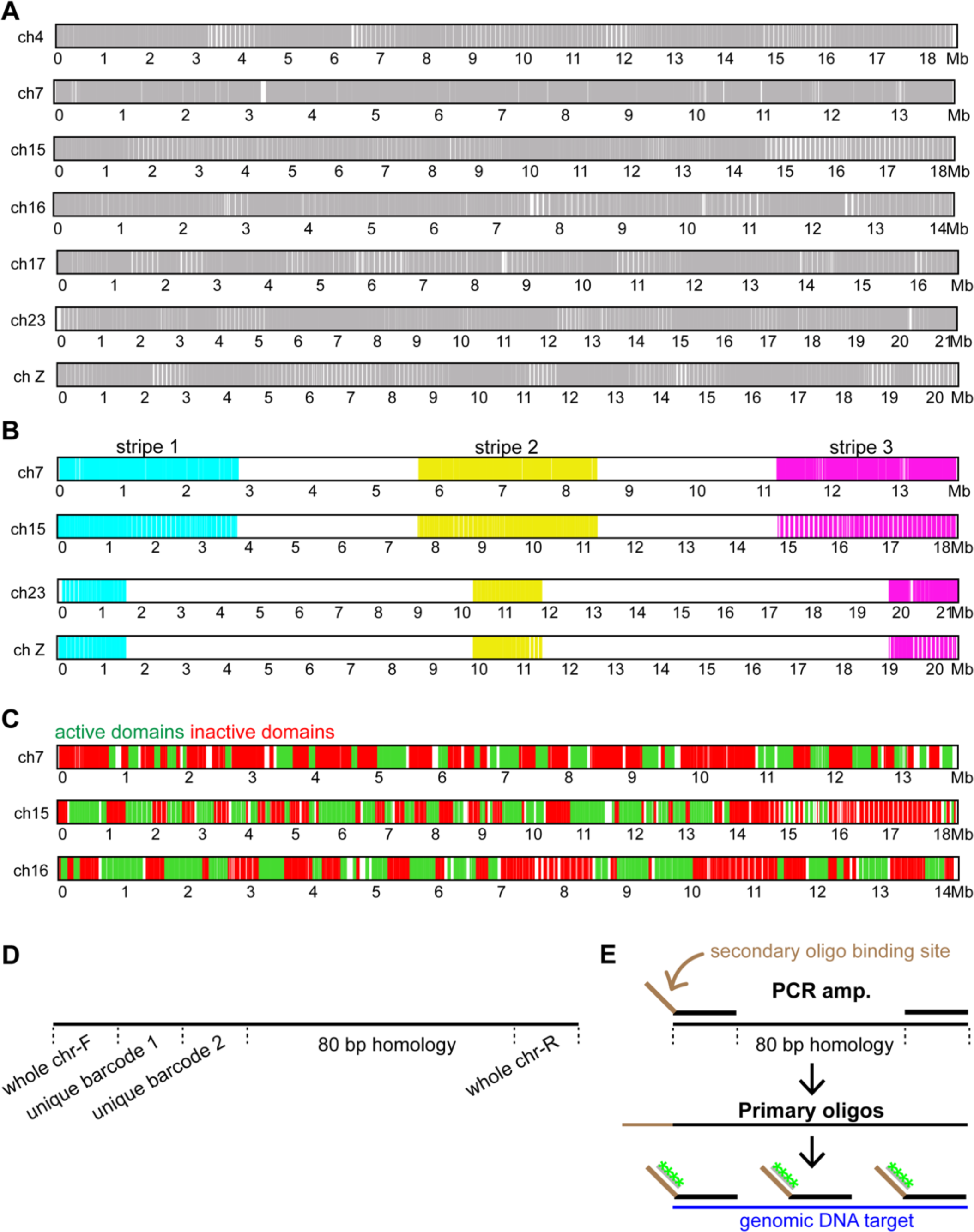
*B. mori* Oligopaint design. A-C) Schematic of Oligopaints in *B. mori*. Whole chromosome Oligopaints are shown in A, stripe Oligopaints in B, and active/inactive Oligopaints in C. White regions indicate the absence of oligos (A) or regions not labeled by the respective barcode indices (B, C). D) Schematic of primary probe design, showing whole chromosome barcodes and two unique barcodes (for stripes or active/inactive domains). E) Schematic for Oligopaint DNA FISH assay with labeled secondary oligos. First, ordered oligos are amplified with primers containing barcode of interest and secondary oligo binding site, generating primary oligos. Primary oligos are then annealed to DNA and labeled with secondary oligos (shown in green).

**Table 1.** Chromosome and chromosome paint information.

| Chrom. | chromosome size (bp) | size painted (bp) | paint start | paint stop | density (probes/kb) | total # of oligos |
| --- | --- | --- | --- | --- | --- | --- |
| 4 | 18737234 | 18639239 | 282 | 18639521 | 1.5 | 26841 |
| 7 | 13944894 | 13868845 | 35931 | 13904776 | 3 | 42625 |
| 15 | 18440292 | 18354755 | 21089 | 18375844 | 1 | 17756 |
| 16 | 14337292 | 14275583 | 27737 | 14303320 | 1.5 | 20190 |
| 17 | 16840672 | 16806551 | 9415 | 16815966 | 1.5 | 23834 |
| 23 | 21465692 | 21339065 | 123188 | 21462253 | 1.5 | 30506 |
| Z (1) | 20666287 | 20578020 | 37936 | 20615956 | 1.5 | 29784 |

### *B. mori* Oligopaints are highly specific

As the *B. mori* genome used to design the Oligopaints is in a semi-draft state (assembled into chromosomes but still with many unmapped contigs), we first tested the specificity of our *B. mori* chromosome paints using karyotype analysis. To this end, meiotic chromosome spreads were prepared from late 4^th^ or early 5^th^ larval testes and ovaries. As silkmoths have a very short adult lifespan (only 5-7 days), meiosis begins early in the larval stages (62). Due to the small, holocentric nature of silkworm chromosomes, mitotic chromosomes are small and highly compact, while chromosomes in meiotic prophase I (pachytene sub-stage; Figure 2A; reviewed in (63)) are more linear due to synapsis (Figure S2)(64), making meiotic chromosomes better suited for our karyotype analyses. Since homologs are paired during most sub-stages of meiotic prophase I (Figure 2A), the expectation was a single fluorescence signal per chromosome. A detailed description of our pachytene chromosome spread protocol used for Oligopaint FISH can be found in the Materials and Methods.

**Figure 2.**
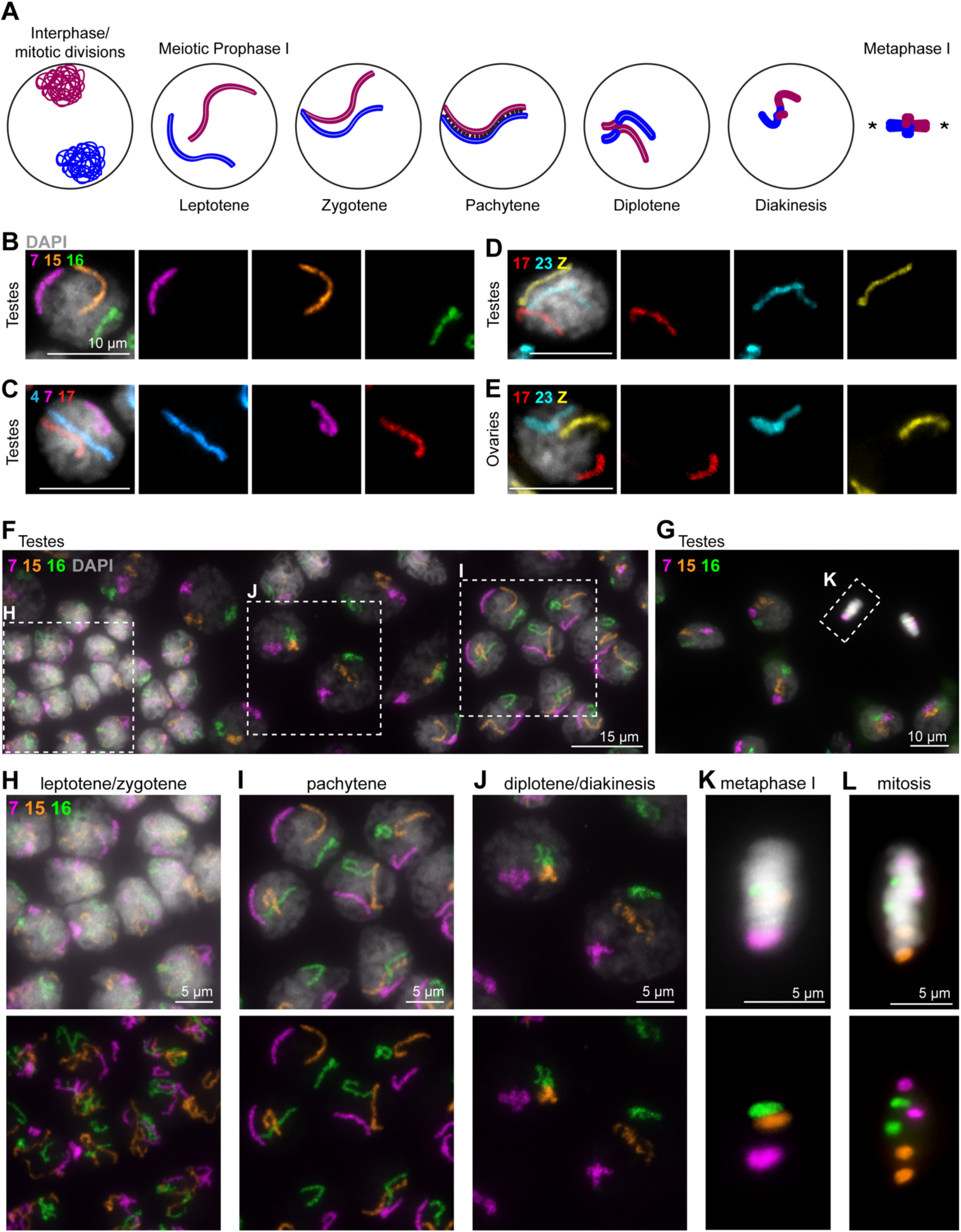
Whole chromosome Oligopaints in *B. mori* 5^th^ instar germline squashes. A) Schematic of early meiosis I (prophase I and metaphase I). One pair of homologous chromosomes is shown (red = paternal; blue = maternal). Prophase I is typically subdivided into five distinct stages: leptotene, zygotene, pachytene, diplotene, and diakinesis. Briefly: *in leptotene, replicated chromosomes are reorganized and compacted into a linear scaffold structure. In zygotene, synapsis begins between the homologous chromosomes. In pachytene, synapsis is complete (black dots represent the synaptonemal complex holding the homologs together). This is also when crossing over can occur. In diplotene, the homologs repulse, condense further, and the SC breaks down. The homologs remain attached via chiasma (crossovers). Finally, in diakinesis, chromosome condensation and cruciform bivalent formation is nearly complete as the cell prepares for metaphase I.* Asters in metaphase I schematic indicate the spindle poles. B-E) Pachytene cells labeled with three whole chromosome Oligopaints, as labeled. A-C, larval testes. D, larval ovary. Scale bars = 10 µm. DAPI is shown in gray. F-G) Meiotic prophase I and metaphase I (F) cells from larval testes squash with whole chromosome paints for ch7 (magenta), ch15 (orange), and ch16 (green). Boxes indicate subsequent panels as indicated. DAPI is shown in gray. H-K) Zooms from E-F, as indicated below. H) Leptotene/zygotene cells, with unpaired, decondensed chromosomes. I) Pachytene cells, with paired, linear, and relatively decondensed chromosomes. J) Diplotene/diakinesis cells, with paired, less linear and more compact chromosomes. K) Metaphase I cells, with paired homologs condensed and aligned along the metaphase plate. L) Mitotic cells from larval testes, with chromosomes condensed and aligned along the metaphase plate but with unpaired homologs.

Using our whole chromosome paints, three chromosomes at a time were labeled on pachytene chromosome spreads from testes and ovaries. This, indeed, illustrated singular and distinct fluorescence signals for each tested chromosome during pachytene (Figure 2B-2E, 2F, and 2I). In our larval testes squashes, we also were able to identify cells in early meiotic prophase I before pairing has completely occurred (leptotene/zygotene; (Figure 2F, 2H). Interestingly, a wide variety of partially paired chromosome configurations were observed at this stage, many of which contain large chromosome loops (Figure S3A). Furthermore, we noticed that all chromosomes do not pair simultaneously, as many cells harbor some paired and some unpaired chromosomes (Figure S3B).

In addition to early prophase, cells in late prophase (post-pachytene) could also be distinguished. In our testes squashes, we identified cells with more condensed, paired chromosomes as being in diplotene/diakinesis (also known as the diffuse stage based on chromatin morphology, Figure 2F, 2J). Furthermore, we were able to identify cells in which chromosomal bivalents are compacted and aligned at the metaphase plate (metaphase I, Figure 2G, 2K). Finally, we observed somatic cells in our testes squahses harboring two distinct fluorescence signals per chromosomes, indicating that homologs are unpaired in the majority of somatic cells in *B. mori* (Figure S4).

In larval ovary squashes, we were able to identify linear, paired pachytene chromosomes (Figure 2E). Additionally, post-pachytene nurse cells could be identified by their unpaired, condensed chromosomes (Figure S5; (58)). Finally, mitotic cells are also present in the gonads, and thus were also used as additional validation of our probe specificity (Figure 2L, S2, and S6). Interestingly, when we tested these same whole chromosome paints on mitotic chromosome spreads from a *B. mori* ovary-derived cell line, BmN4, we found that our probes partially labeled multiple chromosomes, suggesting that the karyotype in these cells has undergone dramatic rearrangements compared to the genome-derived strain (Figure S7). Together, these data not only illustrate that our Oligopaint libraries are specific but also act as a validation for the *B. mori* genome assembly.

**Figure 6.**
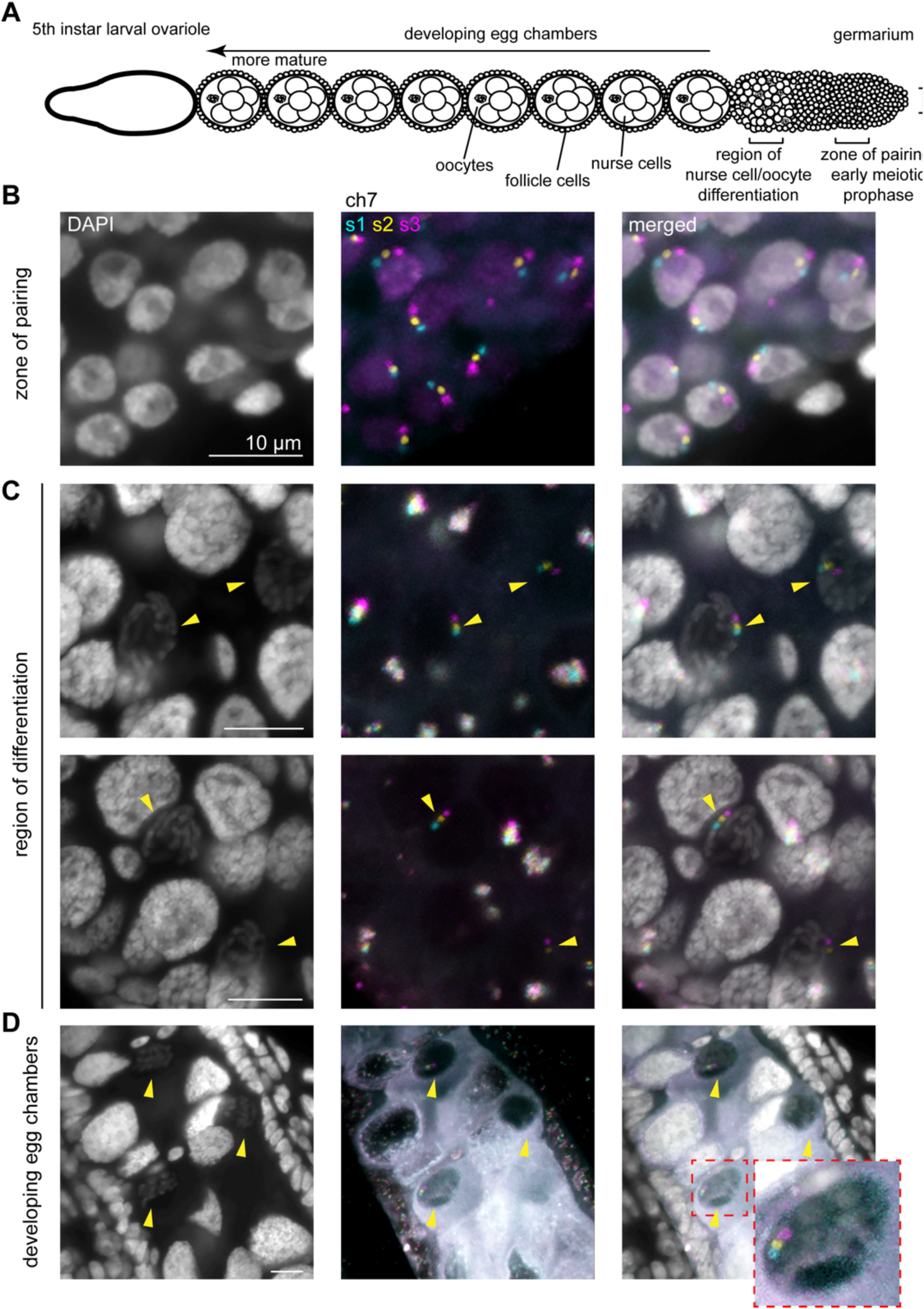
FISH with stripe chromosome paints in whole mount 5^th^ instar larval ovary. A) Schematic of ovariole from 5^th^ instar *B. mori* larval ovary. B) Representative pachytene nuclei from germarium of larval ovary labeled with ch7 stripe paints. C) Representative fields showing early oocytes and early nurse cells labeled with ch7 stripe paints. Yellow arrowheads indicate oocytes. Chromosome wide pairing is still present in oocytes at this stage. D) Representative image showing mature (∼stage 5) egg chambers. Yellow arrowheads indicate oocytes. Chromosome wide pairing is still present in oocytes at this late prophase stage.

### Detection of stripe and chromatin state sub-libraries

While the specificity of our whole chromosome paints indicated that the *B. mori* genome is accurately assembled at the chromosome level, we needed to validate the intra-chromosomal genome assembly and in turn, the specificity of our sub-chromosomal paints. For this, we again turned to pachytene chromosome spreads in the male germline, where the linear nature of chromosomes allowed us to verify the linear order of our probes. We started with the stripe sub-libraries, wherein selected chromosomes were sub-divided into 5 stripes approximately 3 Mb in size, or 13 stripes approximately 1.5 Mb in size, depending on the chromosome (Table 2 and Figure 1B). Only the first, middle, and last stripes were labeled with secondary oligos to visualize three stripes along each chromosome (Figure 1B, 3A, 3B; stripe 1, 2 and 3, respectively). FISH with these stripe paints for ch15 in pachytene spreads from larval testes revealed a singular focus for each stripe, with stripes 1, 2, and 3 positioned in the predicted order along the linear chromosome (Figure 3A). This was also true for ch23 and the Z chromosome (Figure 3B-C).

**Figure 3.**
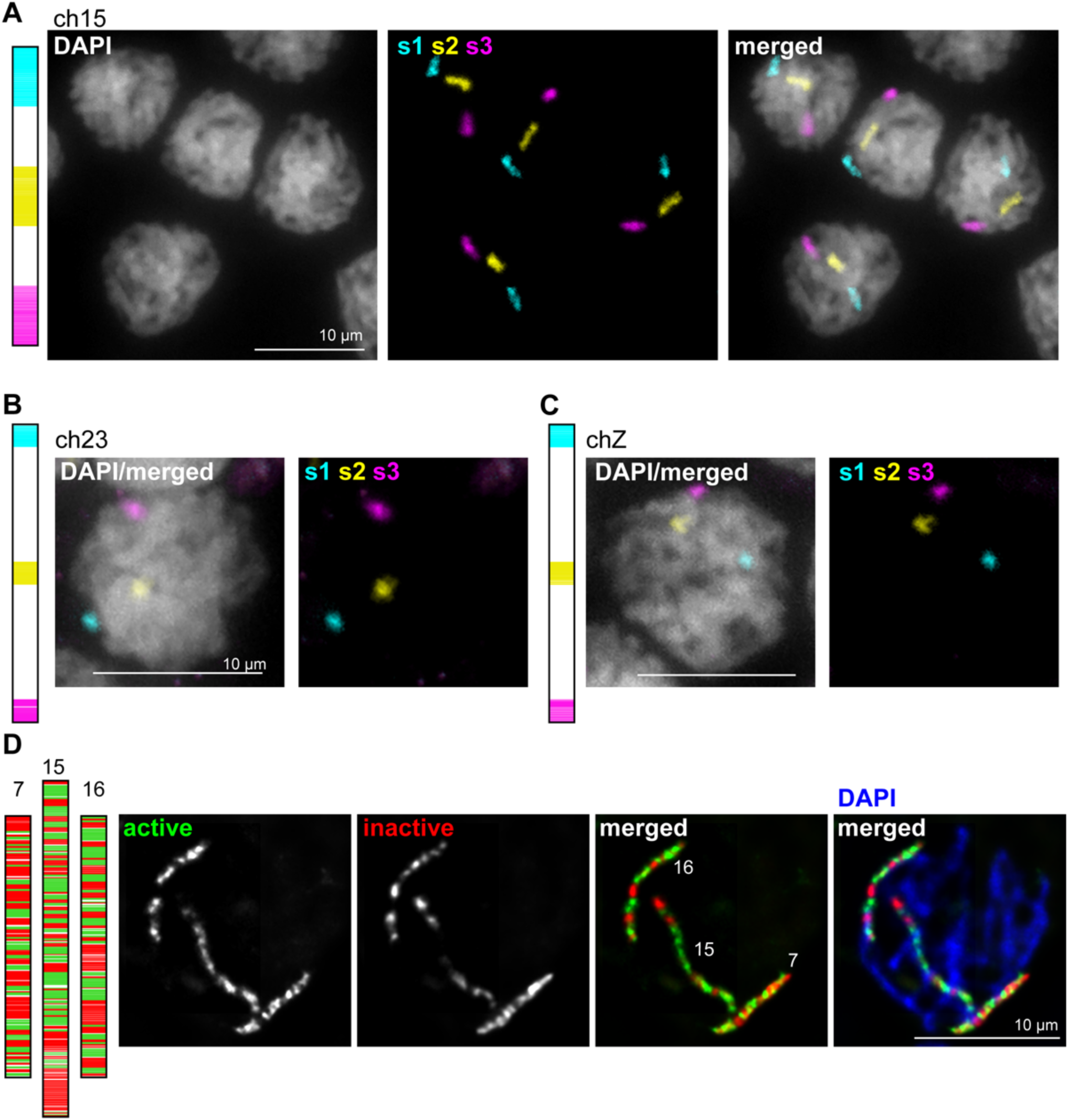
Stripe and active/inactive chromosome paints in *B. mori* 5^th^ instar testes squashes. A) Left: Schematic of stripe paints for ch15, with stripe 1 (s1) in cyan, stripe 2 (s2) in yellow, and stripe 3 (s3) in magenta. Right: Pachytene cells labeled with ch15 stripe paints. B-C) Left: Schematic of stripe paints for ch23 or chZ. Right: Representative pachytene nucleus labeled with stripe paints for ch23 or chZ. D) Left: Schematic of active/inactive paints for ch7, 15, and 16. Active domains are shown in green and inactive domains are shown in red. Right: Representative pachytene nucleus labeled with paints for all 3 chromosomes.

**Table 2.** Stripe sub-library paint information.

| Chrom. and stripe | paint start | paint stop | Stripe size (Mb) |
| --- | --- | --- | --- |
| 7 stripe 1 | 35931 | 2809682 | 2.77 |
| 7 stripe 2 | 5587874 | 8357288 | 2.76 |
| 7 stripe 3 | 11131025 | 13904776 | 2.77 |
| 15 stripe 1 | 21089 | 3684775 | 3.66 |
| 15 stripe 2 | 7380341 | 11061106 | 3.68 |
| 15 stripe 3 | 14748412 | 18375844 | 3.62 |
| 23 stripe 1 | 123188 | 1638259 | 1.51 |
| 23 stripe 2 | 9909172 | 11556867 | 1.65 |
| 23 stripe 3 | 19815237 | 21462253 | 1.65 |
| Z stripe 1 | 37936 | 1589202 | 1.55 |
| Z stripe 2 | 9538407 | 11119201 | 1.58 |
| Z stripe 3 | 19078122 | 20615956 | 1.53 |

**Table 3.**
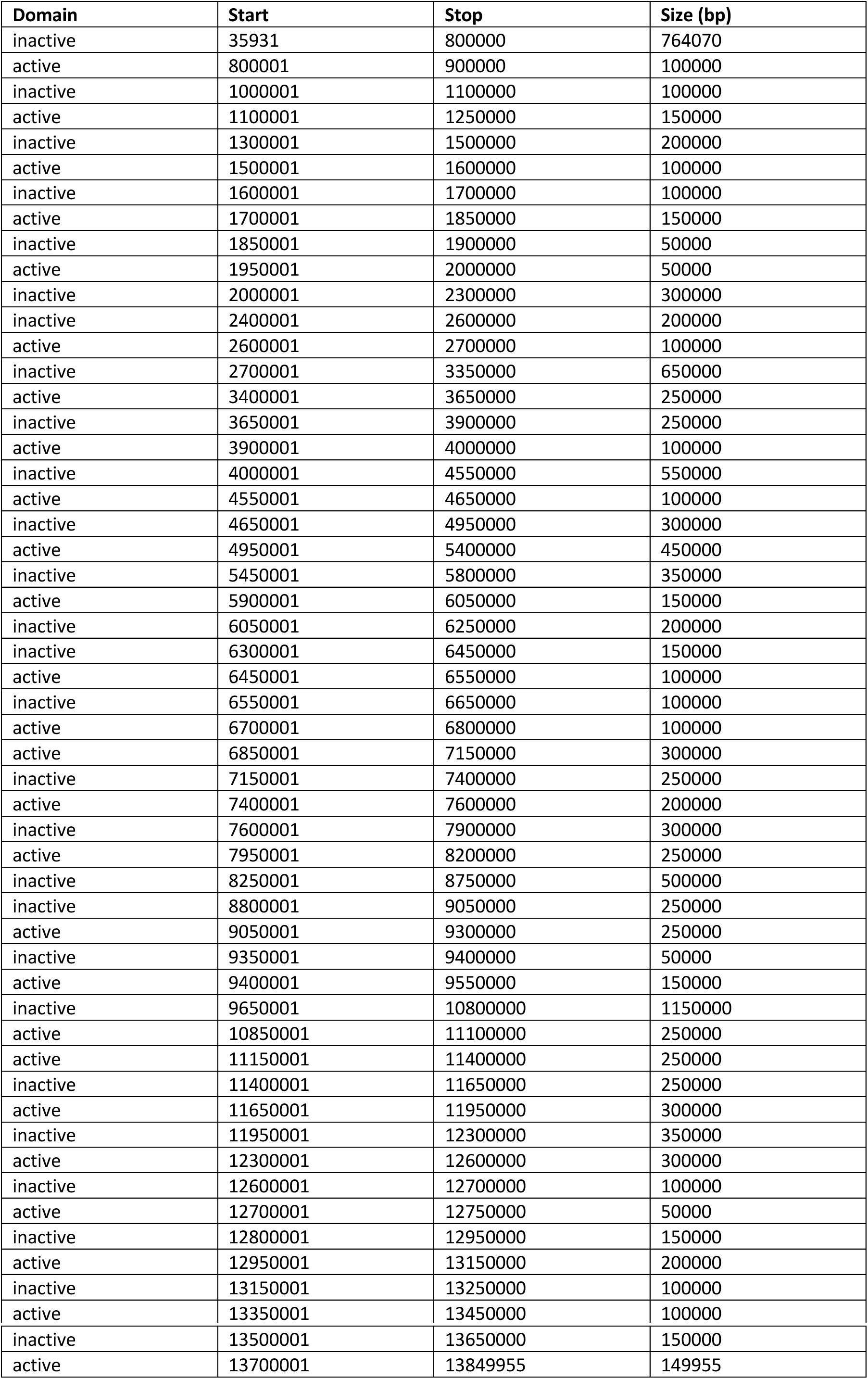
Ch7 Active and inactive chromatin domains paint information.

**Table 4.** Ch15 Active and inactive chromatin domains paint information.

| Domain | Start | Stop | Size (bp) |
| --- | --- | --- | --- |
| inactive | 21089 | 200000 | 178912 |
| active | 250001 | 850000 | 600000 |
| active | 900001 | 1000000 | 100000 |
| inactive | 1000001 | 1400000 | 400000 |
| active | 1400001 | 1950000 | 550000 |
| inactive | 1950001 | 2250000 | 300000 |
| active | 2250001 | 2550000 | 300000 |
| inactive | 2550001 | 2800000 | 250000 |
| active | 2850001 | 3200000 | 350000 |
| inactive | 3200001 | 3500000 | 300000 |
| active | 3550001 | 3850000 | 300000 |
| inactive | 3850001 | 3900000 | 50000 |
| active | 3950001 | 4100000 | 150000 |
| inactive | 4100001 | 4250000 | 150000 |
| active | 4250001 | 4350000 | 100000 |
| inactive | 4350001 | 4650000 | 300000 |
| active | 4650001 | 4750000 | 100000 |
| inactive | 4750001 | 4950000 | 200000 |
| inactive | 5000001 | 5150000 | 150000 |
| active | 5150001 | 5250000 | 100000 |
| inactive | 5300001 | 5350000 | 50000 |
| active | 5350001 | 6100000 | 750000 |
| inactive | 6100001 | 6200000 | 100000 |
| active | 6250001 | 6500000 | 250000 |
| inactive | 6500001 | 6700000 | 200000 |
| active | 6700001 | 6850000 | 150000 |
| inactive | 6900001 | 7100000 | 200000 |
| active | 7100001 | 7300000 | 200000 |
| inactive | 7300001 | 7550000 | 250000 |
| active | 7550001 | 7800000 | 250000 |
| inactive | 7800001 | 8100000 | 300000 |
| active | 8100001 | 8300000 | 200000 |
| inactive | 8400001 | 8650000 | 250000 |
| active | 8650001 | 8750000 | 100000 |
| inactive | 8750001 | 8800000 | 50000 |
| active | 8850001 | 8900000 | 50000 |
| inactive | 8900001 | 9100000 | 200000 |
| active | 9100001 | 9500000 | 400000 |
| inactive | 9500001 | 9550000 | 50000 |
| active | 9600001 | 9750000 | 150000 |
| active | 9800001 | 9950000 | 150000 |
| inactive | 10000001 | 10500000 | 500000 |
| active | 10500001 | 11200000 | 700000 |
| active | 11350001 | 11400000 | 50000 |
| inactive | 11400001 | 11500000 | 100000 |
| active | 11500001 | 11950000 | 450000 |
| inactive | 11950001 | 12000000 | 50000 |
| active | 12050001 | 12200000 | 150000 |
| inactive | 12200001 | 12250000 | 50000 |
| active | 12300001 | 12500000 | 200000 |
| inactive | 12500001 | 12600000 | 100000 |
| active | 12600001 | 12850000 | 250000 |
| inactive | 12850001 | 12950000 | 100000 |
| active | 12950001 | 13400000 | 450000 |
| inactive | 13450001 | 13600000 | 150000 |
| active | 13650001 | 13750000 | 100000 |
| inactive | 13750001 | 14150000 | 400000 |
| inactive | 14200001 | 14850000 | 650000 |
| active | 14900001 | 15050000 | 150000 |
| inactive | 15100001 | 15250000 | 150000 |
| active | 15300001 | 15500000 | 200000 |
| inactive | 15550001 | 15600000 | 50000 |
| active | 15600001 | 15750000 | 150000 |
| inactive | 15750001 | 16000000 | 250000 |
| inactive | 16050001 | 16350000 | 300000 |
| inactive | 16400001 | 17850000 | 1450000 |
| inactive | 17900001 | 18100000 | 200000 |
| active | 18150001 | 18200000 | 50000 |
| inactive | 18200001 | 18250000 | 50000 |
| active | 18300001 | 18350000 | 50000 |
| inactive | 18350001 | 18375844 | 25844 |

**Table 5.** Ch16 Active and inactive chromatin domains paint information.

| Domain | Start | Stop | Size (bp) |
| --- | --- | --- | --- |
| inactive | 27737 | 50000 | 22264 |
| active | 50001 | 150000 | 100000 |
| inactive | 150001 | 250000 | 100000 |
| active | 250001 | 350000 | 100000 |
| inactive | 350001 | 650000 | 300000 |
| active | 700001 | 1350000 | 650000 |
| inactive | 1400001 | 1700000 | 300000 |
| active | 1700001 | 2300000 | 600000 |
| inactive | 2300001 | 2400000 | 100000 |
| active | 2400001 | 2700000 | 300000 |
| inactive | 2700001 | 3200000 | 500000 |
| active | 3200001 | 3600000 | 400000 |
| inactive | 3600001 | 4050000 | 450000 |
| active | 4050001 | 4200000 | 150000 |
| inactive | 4200001 | 4500000 | 300000 |
| active | 4500001 | 4600000 | 100000 |
| active | 4700001 | 4750000 | 50000 |
| inactive | 4750001 | 4800000 | 50000 |
| active | 4900001 | 4950000 | 50000 |
| active | 5000001 | 5250000 | 250000 |
| inactive | 5250001 | 5600000 | 350000 |
| active | 5600001 | 6000000 | 400000 |
| inactive | 6000001 | 6150000 | 150000 |
| active | 6200001 | 6250000 | 50000 |
| active | 6300001 | 6500000 | 200000 |
| active | 6550001 | 6650000 | 100000 |
| inactive | 6650001 | 6850000 | 200000 |
| active | 6850001 | 7000000 | 150000 |
| inactive | 7050001 | 7950000 | 900000 |
| inactive | 8000001 | 8500000 | 500000 |
| active | 8550001 | 8750000 | 200000 |
| inactive | 8800001 | 8900000 | 100000 |
| active | 8900001 | 9400000 | 500000 |
| inactive | 9400001 | 9450000 | 50000 |
| active | 9500001 | 10100000 | 600000 |
| inactive | 10100001 | 11450000 | 1350000 |
| active | 11450001 | 11550000 | 100000 |
| inactive | 11550001 | 11900000 | 350000 |
| active | 11950001 | 12250000 | 300000 |
| inactive | 12250001 | 12400000 | 150000 |
| active | 12400001 | 12500000 | 100000 |
| inactive | 12500001 | 12700000 | 200000 |
| active | 12750001 | 13200000 | 450000 |
| inactive | 13250001 | 13800000 | 550000 |
| active | 13800001 | 14100000 | 300000 |
| inactive | 14100001 | 14303320 | 203320 |

To test the specificity of our transcriptionally active and inactive chromatin domain paints, we labeled all three chromosomes with this barcode index (ch7, 15, and 16) at the same time on larval testes pachytene spreads. This revealed three separate linear chromosomes with distinct banding patterns (Figure 3D) corresponding to the respective paint schematic (Figure 1 and 3D). Together, these results indicate that the intra-chromosomal assembly for these chromosomes is highly accurate and our paints are specific for their target chromosomal domains.

### Telomeres face poleward at random in metaphase I bivalents in testes

In addition to identifying pachytene cells in our assays with the stripe paints in larval testes, we were able to identify cells in all stages of meiosis I up to metaphase I, as well as cells in interphase (Figure 4A). Interestingly, we noted that traditional cruciform bivalents are formed at metaphase I, and these bivalents are highly reminiscent of those seen in meiosis in the nematode *C. elegans*, with one telomere remaining paired and the other telomere facing poleward ((Figure 4B-D) reviewed in (44)). While this “telokinetic” chromosome configuration was previously hypothesized to occur in meiosis in *B. mori*, it has never before been directly observed.

**Figure 4.**
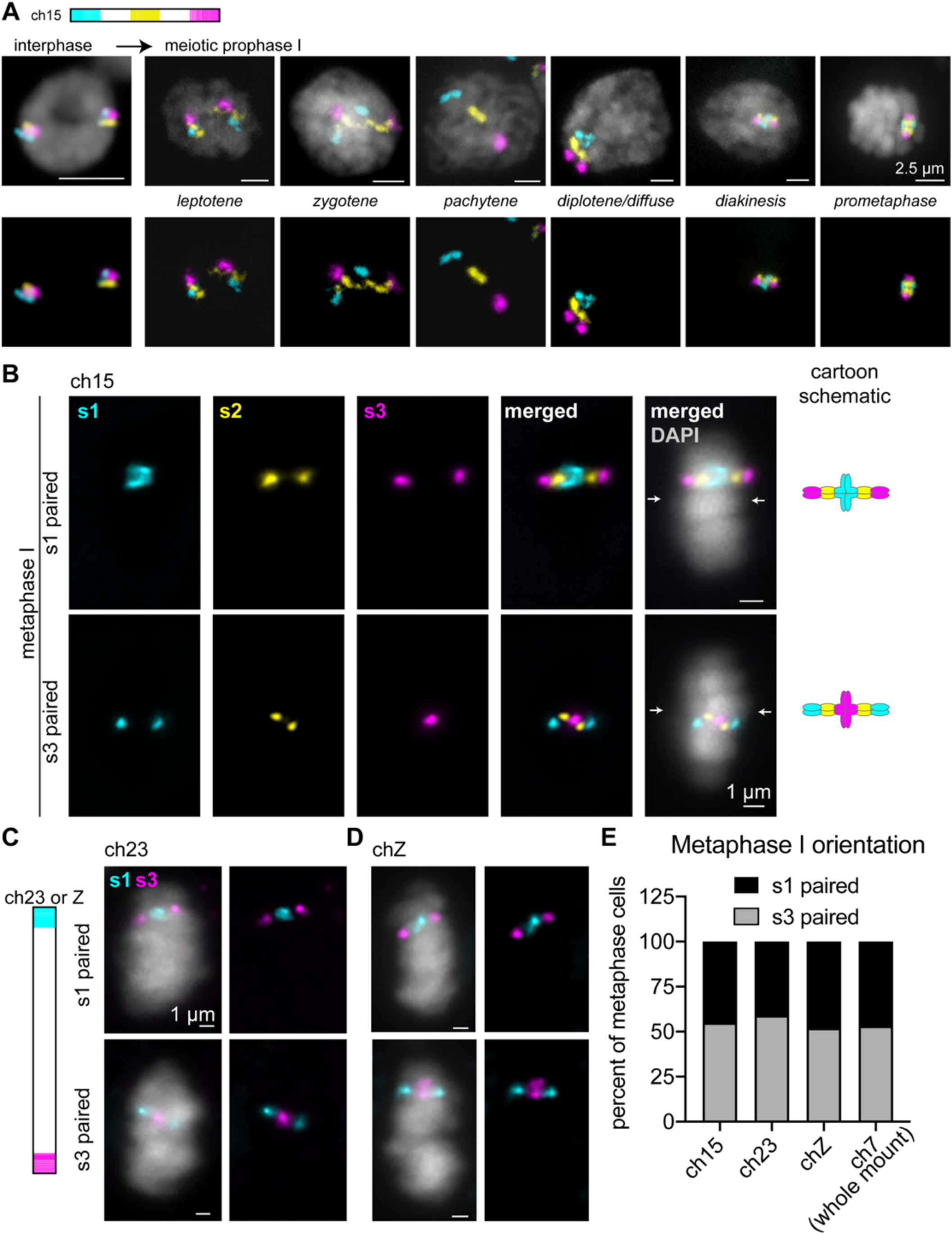
Analysis of pairing and metaphase I bivalent formation in 5^th^ instar larval testes squashes. A) Left: Schematic of stripe paints for ch15, with stripe 1 (s1) in cyan, stripe 2 (s2) in yellow, and stripe 3 (s3) in magenta. Right: representative nuclei at the designated stages labeled with ch15 stripe paints. When cells enter meiosis, chromosomes begin to decondense (leptotene) and homologs pair (zygotene). Pairing is complete by pachytene, with complete synapsis for crossing over, and chromosomes are linear. Chromosomes begin to condense for segregation in diplotene and diakinesis. DAPI is shown in gray. B) Metaphase I bivalents labeled with ch15 stripe paints. Top: bivalent with pairing in stripe 1 domain. Bottom: bivalent with pairing in stripe 3 domain. Schematics of bivalents shown on the right. C) Left: Schematic of stripe paints for ch23 or Z, showing only stripe 1 (cyan) and stripe 3 (magenta). Right: representative metaphase I cells showing pairing in the s1 domain (top) or s3 domains (bottom) for ch23. D) Representative metaphase I cells showing pairing in the s1 domain (top) or s3 domains (bottom) for chZ. E) Quantification of metaphase I orientation for ch15, 23, Z, and 7 (from whole-mount larval testes). Ch15, n=182 (45% s1 paired). Ch23, n=144 (41% s1 paired). ChZ, n=169 (48% s1 paired). Ch7, n=96 (47% s1 paired). Each FISH assay was performed on a different larva.

We next wanted to determine if both telomeres of a chromosome can act as localized kinetochores during meiosis in *B. mori* or if one telomere preferentially faces poleward. In *C. elegans*, either telomere on any given chromosome can harbor kinetochore activity, and both do so at random depending on where crossovers form during meiotic prophase I (42, 49). To test whether a similar mechanism occurs in *B. mori*, we examined metaphase I bivalents in larval testes using our stripe paints for chromosomes 15, 23, and Z. Quantification of which telomere remains paired and which telomere faces poleward (stripe 1 or stripe 3) revealed that approximately half of metaphase I cells harbor pairing in the stripe 1 domain and half harboring pairing in the stripe 3 domain for all tested chromosomes, including ch Z (Figure 4E). This finding suggests, like nematodes, *B. mori* telomere regions likely act as localized kinetochores during meiosis. Furthermore, the orientation of chromosomes in the bivalent is not pre-determined, with both telomeres having an equal probability of being either paired or facing poleward.

To further validate this finding, we repeated the experiment using chromosome 7 stripe paints in whole mount late 5^th^ instar larval testes (Figure S8). As we predicted based on our testes squashes and previous studies (54), 5^th^ instar larval testes harbor mitotic cells with unpaired homologs that highly resemble mitotic cells seen in whole-mount embryos (Figure 5A-B; Figure S9) and primary spermatocytes at all stages of meiosis I (Figure 5A-D). Interestingly, *B. mori* and other Lepidopteran insects utilize two distinct spermatogenic pathways, ultimately resulting in apyrene (without nuclei) and eupyrene (with nuclei) sperm (65–71). In whole-mount testes, we were clearly able to identify eupyrene secondary spermatocyte bundles (Figure 5C-3) and mature eupyrene sperm (Figures S10 and S11). Additionally, we identified secondary spermatocyte bundles appearing to be apyrene- destined, where some cells have no DNA and the FISH signal is instead diffuse in the cytoplasm, suggesting that the cells in these bundles are beginning the process of nuclear degradation (Figure 5C-4). Importantly, quantification of metaphase I bivalent formation in whole mount testes was completely in agreement with our findings from squashes, showing ch7 stripe 1 paired in 46.9% of cells and ch7 stripe 3 paired in 53.1% (Figure 4E). Together, these findings suggest that *B. mori* chromosomes form traditional bivalent structures at metaphase I with localized centromere activity restricted to one telomeric region at random.

**Figure 5.**
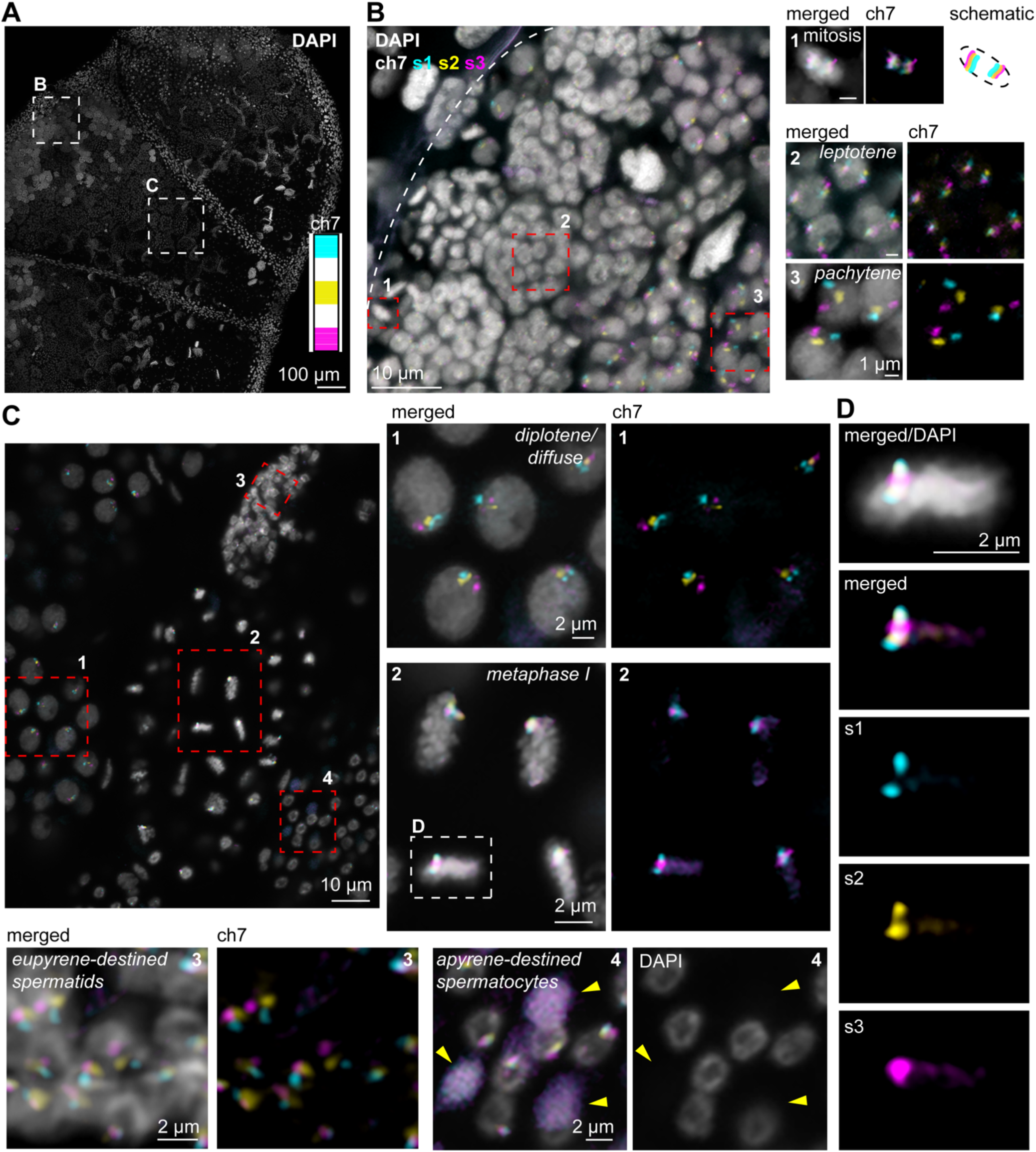
FISH with stripe chromosome paints in whole mount 5^th^ instar larval testes. A) 5^th^ instar larval testis stained with DAPI. Boxes indicate subsequent panels as indicated. Inset: ch7 stripe paints used in B-D. B) Zoom of mitotic and pachytene region of larval testes as shown in A, labeled with ch7 stripe paints. Red boxes indicate zooms shown to the right. 1) mitotic cells – chromosomes condensed, homologs unpaired, and aligned at the metaphase plate. Note how chromosomes are compacted perpendicular to the metaphase plate. 2) leptotene cells, chromosomes are slightly decondensed and homologs are unpaired. 3) pachytene cells, chromosomes are paired head-to-tail and linear. C) Zoom of late prophase/metaphase I region of larval testis as shown in A, labeled with ch7 stripe paints. Red boxes indicate zooms shown to the right and below. 1) diplotene cells (diffuse stage), chromosomes are still paired and beginning to condense. Left: merged with DAPI in gray. Right: ch7 stripe 1 in cyan, stripe 2 in yellow, and stripe 3 in magenta. 2) metaphase I cells labeled with ch7 stripe paints. White box indicates zoom shown in D. Left: merged with DAPI in gray. Right: stripe 1 (cyan) and stripe 3 (magenta) paints. 3) eupyrene-destined secondary spermatocyte bundle. Left: merged ch7 stripe paints with DAPI in gray. Right: ch7 stripe paints, stripe 1 (cyan) and stripe 3 (magenta). 4) apyrene-destined secondary spermatocyte bundle. Left: merged ch7 stripe paints with DAPI in gray. Right: DAPI. Yellow arrowheads indicate spermatocytes that have already undergone nuclear degradation. Scale bar = 2 um for all panels 1-4. D) Zoom of metaphase I cell indicated in C-2. Top: merged with DAPI in gray. Bottom: ch7 stripe 1 (cyan), stripe 2 (yellow), and stripe 3 (magenta) paints. Bivalent pairing is in stripe 3 domain in this cell. Note: zoomed fields for all panels may display a slightly different Z position than the larger field views.

### Chromosome-wide homolog pairing is persistent in *B. mori* female larval ovaries

In contrast to what we and others have observed in *B. mori* males, homolog pairing in *B. mori* females is reported to be unconventional, without chiasma formation and with the SC transforming into “elimination chromatin” over one micron in width (54,57–59). Despite these previous observations, we found that a significant number of nuclei with homologs entirely paired in meiotic chromosome spreads from late 4^th^/early 5^th^ instar larvae (Figure 2 and S5). This finding led us to wonder whether homolog pairing is more stable in *B. mori* female meiosis than previously appreciated. Like *Drosophila*, the *B. mori* larval ovary is composed of polytrophic meroistic ovarioles containing linear arrays of developing egg chambers, with the tip (germarium) harboring germ line stem cells that are mitotically dividing and the most mature chambers being the most distal from the stem cell niche ((62,72,73), Figure 6A and Figure S12). However, as moths have a much shorter adult lifespan than flies (only 5-7 days for silkmoths versus 2-3 months for *Drosophila*), the majority of oogenesis occurs in the larval and pupal stages (62).

To determine how long homologs remain paired end-to-end in meiotic prophase in females, we performed whole-mount DNA FISH with ch7 stripe paints in 5^th^ instar larval ovaries. This approach yielded robust FISH signal throughout the developing egg chambers and in the germarium (Figure 6). In agreement with our initial observations from ovary squashes, we observed that all cells in the germarium harbor paired, linear homologs through early pachytene (Figure 6A-B). Interestingly, ch7 homologs were still paired end-to-end in late pachytene, where nurse cells and the developing oocytes begin to differentiate (Figure 6C). The majority of oocytes in the developing egg chambers outside the germarium in 5^th^ instar larvae are arrested in late diakinesis or metaphase I (57), which should be after transformation of the SC at the end of pachytene. Surprisingly, we observed that chromosome-wide pairing persists in all oocytes present in developing egg chambers throughout the length of the 5^th^ instar larval ovarioles (Figure 6C-D). This finding suggests that, even in the absence of chiasma and with the partial breakdown and transformation of the SC, end-to-end pairing persists throughout meiotic prophase I in female *B. mori*. Altogether, these studies using Oligopaints in *B. mori* larval ovaries and testes demonstrate that this FISH-based approach is highly successful in both squashed and whole tissue and can be used to study chromosome dynamics throughout different stages of meiosis.

## Discussion

The silkworm, *B. mori*, has been a model system for studying meiotic chromosomes for decades. Like *Drosophila*, silkworms are readily reared in a laboratory setting and highly amenable to genetic manipulations including RNAi and CRISPR. However, unlike fruit flies, silkworms are large in size, combining the increased ease of dissection and structure visualization commonly associated with mammals with the short generation time of an insect system. Additionally, the large amount of tissue provided by *B. mori* increases the feasibility of genomics assays and other cell population-based approaches, which require a large number of cells. Importantly, the recent sequencing of the *B. mori* genome revealed that there is a high degree of sequence homology between silkworm genes and mammalian disease genes (61,74–76). Furthermore, *Bombyx* represents an excellent insect model system for studying meiosis, as SC constituents in non-Drosophilid arthropods are closely aligned with vertebrates, while *Drosophila* harbor a unique suite of SC factors (77, 78). Finally, *B. mori* harbor 28 chromosomes while humans have 23 (and *Drosophila* have only 4), and our studies along with others have illustrated that homologs remain unpaired in *B. mori* somatic cells (79). This is in stark contrast to the high levels of somatic homolog pairing seen in *Drosophila* (80, 81), making the study of *B. mori* genome dynamics more directly relevant to human biology.

Here, we take advantage of the *B. mori* model system to visualize single chromosome dynamics in meiosis using the Oligopaints technology. As previously described for *C. elegans*, we have utilized the flexibility and scalability of the Oligopaint design process to add chromosome-specific barcodes to label either whole chromosomes or different sub-chromosomal loci using the same set of oligos (19). Using these multiplexed Oligopaints, we have provided the first extensive characterization of single, whole chromosomes in meiosis in both males and females in any species. Our studies show in great detail how chromosomes in larval testes condense, pair, and partially unpair to form metaphase I cruciform bivalents. While crossing over has been reported in male meiosis in *B. mori* (53, 57), clear chiasmata were not apparent in post-pachytene spermatocytes using our imaging approach (Figure 2, 4, 5). We think this inability to detect chiasmata is likely due to the small size and compact nature of *B. mori* chromosomes in diplotene.

We show that mitotic chromosomes in *B. mori*, which are holocentric in structure, align parallel to the metaphase plate, with both telomeres being aligned with the plate and homologs being unpaired. Our studies further reveal that, like those in *C. elegans*, *B. mori* chromosomes do not retain the holocentric configuration in meiosis. Instead, meiotic chromosomes at metaphase I in spermatogenesis align perpendicular to the metaphase plate such that telomeric regions face the spindle poles and likely act as localized kinetochores. Moreover, we demonstrate that both telomeres are equally likely to face poleward and harbor kinetochore activity. A similar telokinetic approach to meiosis has also been observed in the holocentric milkweed bug *Oncopeltus fasciatus* (82) and the kissing bug *Triatoma infestans* (83). Whether crossover position dictates bivalent structure in *B. mori* or other holocentric insects, as in *C. elegans* (42,46,49–51), remains to be explored. Additionally, how the meiotic spindle attaches to *B. mori* chromosomes and what kinetochore proteins are involved is yet to be determined, although electron microscopy studies have suggested that microtubules directly penetrate the poleward surface of chromosomes during spermatogenesis (52). Interestingly, the broadly conserved centromere-specific histone H3 variant CENP-A is absent from the genome of Lepidopteran insects (56), suggesting that both mitotic and meiotic chromosome segregation in *Bombyx* are independent of CENP-A.

We determine that chromosome-wide pairing in *B. mori* female meiosis is more stable than previously appreciated and persists throughout the entirety of meiotic prophase. How chromosomes remain paired end-to-end, even after loss of the central elements of the SC and through SC transformation, remains unclear. Finally, in addition to demonstrating the feasibility of using Oligopaints to study meiotic chromosomes, our studies illustrate that Oligopaints can be designed for species with draft genome assemblies and that the Oligopaints can in turn be used to validate both inter- and intra-chromosome level genome assemblies.

## Materials and Methods

### *B. mori* strains and cell line

Embryos were obtained from Carolina Biological (Burlington, NC), Coastal Silkworms (Jacksonville, FL), Mulberry Farms (Fallbrook, CA), or were freshly laid in the lab from larvae derived from embryos from these sources. Some larvae were obtained from Rainbow Mealworms (Compton, CA). Embryos were kept at 4°C for less than 1 month. For rearing, embryos were transferred to 28°C and larvae were fed fresh mulberry leaves or powdered mulberry chow (Carolina Biological or Rainbow Mealworms). BmN4 cells are commercially available from ATCC (Manassas, VA).

### Oligopaint design and synthesis

Oligopaint libraries were designed as described in the main text. Active and inactive domains were determined primarily based on CENP-T depletion or enrichment. CENP-T ChIP-seq profiles were obtained from BmN4 cells and domains were called as previously described (37) with the following modifications: CENP-T ChIP-seq signal originally in 10 kb windows was averaged over 50 kb. Subsequently, negative CENP-T domains were subtracted from positive CENP-T domains to obtain final CENP-T depleted domains. As previously observed, domains enriched for CENP-T were shown to strongly correlate with enrichment for the repressive histone mark H3K27me3, while domains depleted of CENP-T were shown to strongly correlate with enrichment of the active chromatin marks H3K4me3 and H3K36me3. All information regarding genomic coordinates for Oligopaints and probe density can be found in Tables 1-5. Oligo pools were purchased from CustomArray/GenScript (Redmond, WA; ch 7, 15, 16) or Twist Biosciences (San Francisco, CA; ch 4, 17, 23, Z). Oligopaints were synthesized as previously described by adding barcodes to each oligo for PCR-based amplification (17,30,84).

### Preparation of meiotic chromosome spreads and DNA FISH

For meiotic squashes, late 4^th^ instar or early 5^th^ instar larvae (approximately 3 inches in length) were sacrificed by decapitation. The caterpillars where then cut open anterior to posterior and fileted on a silicone dissecting dish using standard sewing needles. Gonads were harvested using forceps and placed into 1.5 mL tubes containing SF900 tissue culture media. Gonads were then rinsed thrice in 1X PBS, then incubated in 1X PBS+0.5% sodium citrate for 8-10 min. Using forceps, gonads were then transferred to siliconized coverslips (1 gonad per coverslip) and covered with ∼10 μL of 45% acetic acid/1% PFA/1X PBS and fixed for 6 min. Using a poly-L-lysine coated glass slide, gonads were then physically squashed and slide/coverslip were flash frozen in liquid nitrogen. After carefully removing slides from liquid nitrogen, coverslips were removed with a razor blade, and slides were post-fixed in cold (pre-chilled to -20°C) 3:1 methanol:glacial acetic acid for 10 min. After fixation, slides were washed thrice in 1X PBS and subjected to an ethanol row at -20°C (70%, 90%, 100% ethanol, 5 min each) before drying completely at room temp. Slides were dried for 24-72 h.

FISH on meiotic squashes was performed as previously described for mitotic spreads (31). Briefly, after drying slides, slides were denatured at 72°C for 2.5 min in 2xSSCT/70% formamide before again drying with an ethanol row at -20°C. Slides were then left to air dry for 10 min at room temperature. Primary Oligopaint probes were resuspended in hybridization buffer (10% dextran sulfate/2xSSCT/50% formamide/4% polyvinylsulfonic acid), placed on slides, covered with a coverslip, and sealed with rubber cement. Slides were denatured on a heat block in a water bath set to 92°C for 2.5 min, after which slides were transferred to a humidified chamber and incubated at 37°C overnight. The next day, coverslips were removed using a razor blade and slides were washed as follows: 2×SSCT at 60°C for 15 min, 2×SSCT at RT for 15 min, and 0.2×SSC at RT for 5 min. Fluorescently labeled secondary probes were then added to slides, again resuspended in hybridization buffer, covered with a coverslip, and sealed with rubber cement. Slides were incubated at 37°C for 2 h in a humidified chamber, before repeating the above washes. All slides were stained DAPI and mounted in Prolong Diamond (Invitrogen/ThermoFisher, Waltham, MA). Slides were cured overnight before sealing with clear nail polish and imaging.

### FISH on whole-mount gonads and embryos

For whole-mount DNA FISH in gonads, ovaries and testes from late 5^th^ instar larvae (after secession of eating) were dissected in SF900 cell culture media, washed thrice briefly with 1X PBS, and then fixed for 30 min in 4% PFA in PBS with 0.1% Triton-X-100 (0.1% PBS-T) at RT. Gonads were then washed again thrice in 1X PBS and permeabilized with 0.5% PBS-T for 15 min at RT. Gonads were pre-denatured by washing as follows: 2xSSCT for 10 min at RT, 2xSSCT/20% formamide for 10 min at RT, 2xSSCT/50% formamide for 10 min at RT, 2xSSCT/50% formamide for 3 h at 37°C, 2xSSCT/50% formamide for 3 min at 92°C, 2xSSCT/50% formamide for 20 min at 60°C. To the 2xSSCT/50% formamide, 100 pmol of each probe was directly added. Gonads were then denatured at for 3 min at 92°C and incubated overnight at 37°C. The next day, gonads were washed: 3x 30 min each in 2xSSCT/50% formamide at 37°C, 1x 15 min in 2xSSCT at RT. 20 pmol of each secondary oligo was added with 50% formamide and incubated for 3 h at 37°C. Final washes were performed (2x 30 min in 2xSSCT/50% formamide at 37°C, 1x 10 min in 2xSSCT/50% formamide at RT, 1x 10 min in 2xSSCT/20% formamide at RT, 1x 10 min in 2xSSCT at RT), gonads were stained with DAPI, and mounted on slides with Prolong Diamond (Invitrogen/ThermoFisher).

For whole-mount embryo FISH, diapausing embryos were removed from 4°C and kept at RT for 3-5 d. Chorions were weakened by soaking in 50% bleach for 15 min and then manually removed with forceps. Embryos were subsequently fixed for 30 min in 4% PFA in 0.1% PBS-T at RT, and FISH was performed as described above for whole-mount gonads.

### Meiotic staging

Stages of meiosis were determined based largely on DAPI morphology and/or cell position in whole-mount gonads. Late 4^th^-early 5^th^ instar male larvae were used for squashes as they only possess primary spermatocytes in meiosis I. In whole-mount testes, meiosis I and II were distinguished based on position in the gonad and based on the number of cells per bundle (with meiosis I bundles harboring approximately 64 cells and meiosis II bundles harboring approximately 128 cells).

### Mitotic spreads from BmN4 cells

To induce mitotic arrest, approximately 1 × 10^5^ cells were treated with 0.5 µg/mL Colcemid Solution (Gibco/ThermoFisher) for 2 h in a 28°C heat block. Cells were then spun for 5 min at 600 x *g* at room temperature to pellet and resuspended in hypotonic solution (500 mL of 0.5% sodium citrate). Cells were incubated in hypotonic solution for 8 min. 100 µL of the cell suspension were then placed in a cytofunnel and spun at 1200 rpm for 5 min with high acceleration using a cytocentrifuge (Shandon Cytospin 4; ThermoFisher). For FISH, slides were then fixed in cold 3:1 methanol: acetic acid for 10 min and washed 3 times for 5 min in PBS-T (PBS with 0.1% Triton X-100). FISH was performed as described above for meiotic spreads.

### Imaging, quantification, and data analysis

Images of meiotic squashes were acquired on a Leica DMi6000 wide-field inverted fluorescence microscope using an HCX PL APO 63x/1.40-0.60 Oil objective (Leica Biosystems, Buffalo Grove, IL), Leica DFC9000 sCMOS Monochrome Camera, and LasX software. Whole mount images were acquired using a Zeiss LSM 780 point scanning confocal (Zeiss Microscope Systems, Jena, Germany) with high sensitivity 32 anode Hybrid-GaAsP detectors. BmN4 mitotic spreads were acquired on a Zeiss AxioObserver Z1 wide-field inverted fluorescence microscope with 100x/1.4 oil Plan-APO objective, a Hamamatsu C13440 ORCA-Flash 4.0 V3 Digital CMOS camera, and ZEN blue software. Images were processed using Huygens deconvolution software (SVI, Hilversum, Netherlands), and tiffs were created in ImageJ. Meiotic bivalent quantification was performed manually.

## Acknowledgements

We would like to thank the members of the Lei and Drinnenberg labs, as well as Jamie Walters, Petr Nguyen, and Martina Dalikova for helpful discussion. We would also like to thank Stacie Hughes, Scott Hawley, Jean-René Huynh, and Anahi Molla-Herman for critical reading of the manuscript.

## Funding

This work was funded by the Intramural Program of the National Institute of Diabetes and Digestive and Kidney Diseases, National Institutes of Health (DK015602 to E.P.L.), and by LabEx DEEP (ANR-11-LABX-0044, ANR-10-IDEX-0001-02 to I.A.D.), an ATIP-AVENIR research grant, Institut Curie, the CNRS and the ERC (CENEVO-758757 to I.A.D.). The funders had no role in study design, data collection and analysis, decision to publish, or preparation of the manuscript.

## Competing Interests

The authors declare no competing interests.

## Supplemental Figures

**Figure S1.**
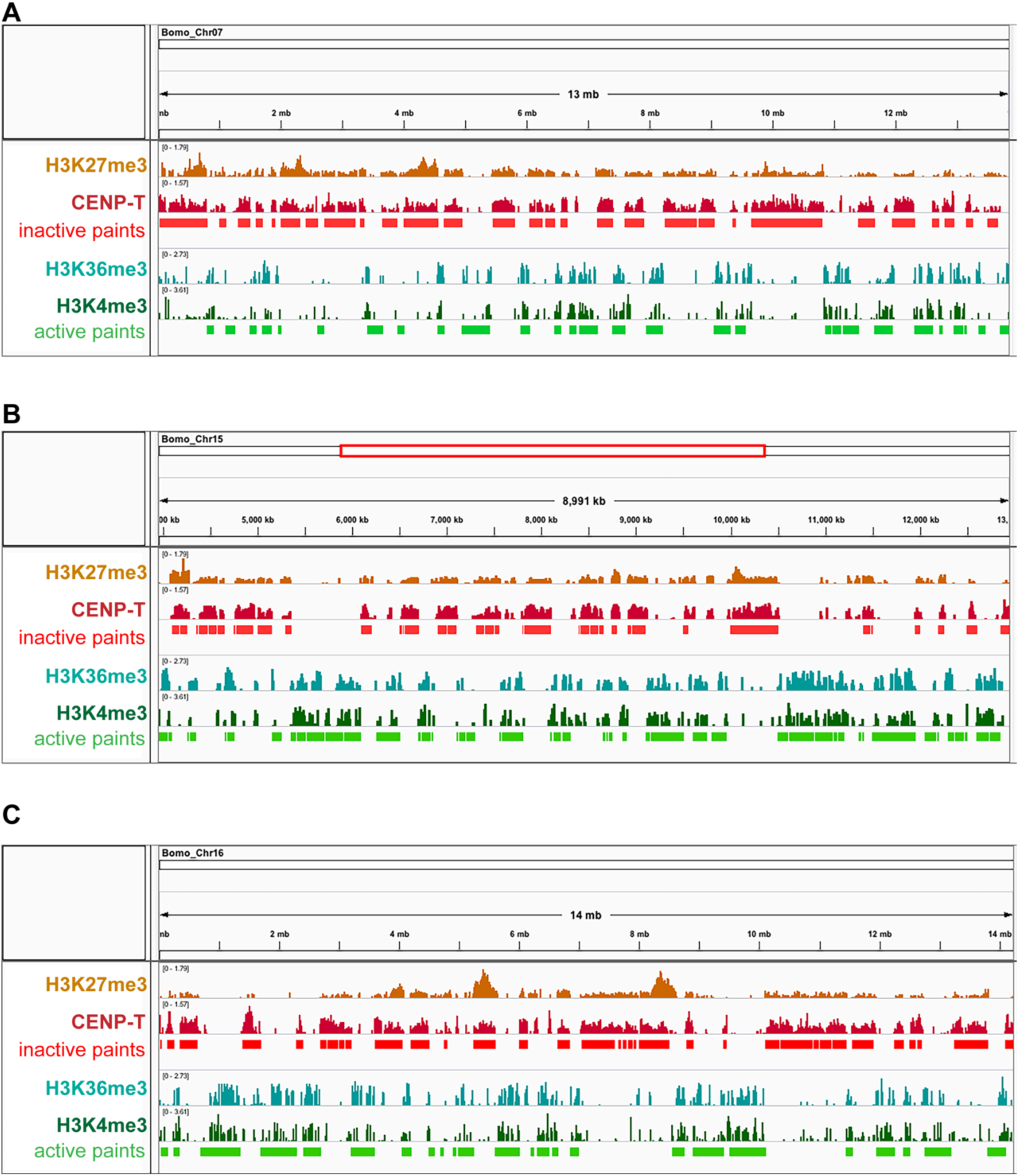
ChIP-seq profiles used to design active and inactive chromosome paints. Screenshots of ChIP-seq data used to design active/inactive chromosome paints. Inactive: H3K27me3 (orange), Centromere Protein T (CENP-T; dark red). Inactive paint domains shown in bright red. Active: H3K36me3 (teal), H3K4me3 (dark green). Active paint domains shown in bright green. ChIP-seq data were previously published (see Materials and Methods). Chromosome 7 is shown in A, part of chromosome 15 is shown in B (coordinates Chr15:3,900,000-12,891,000), and chromosome 16 is shown in C.

**Figure S2.**
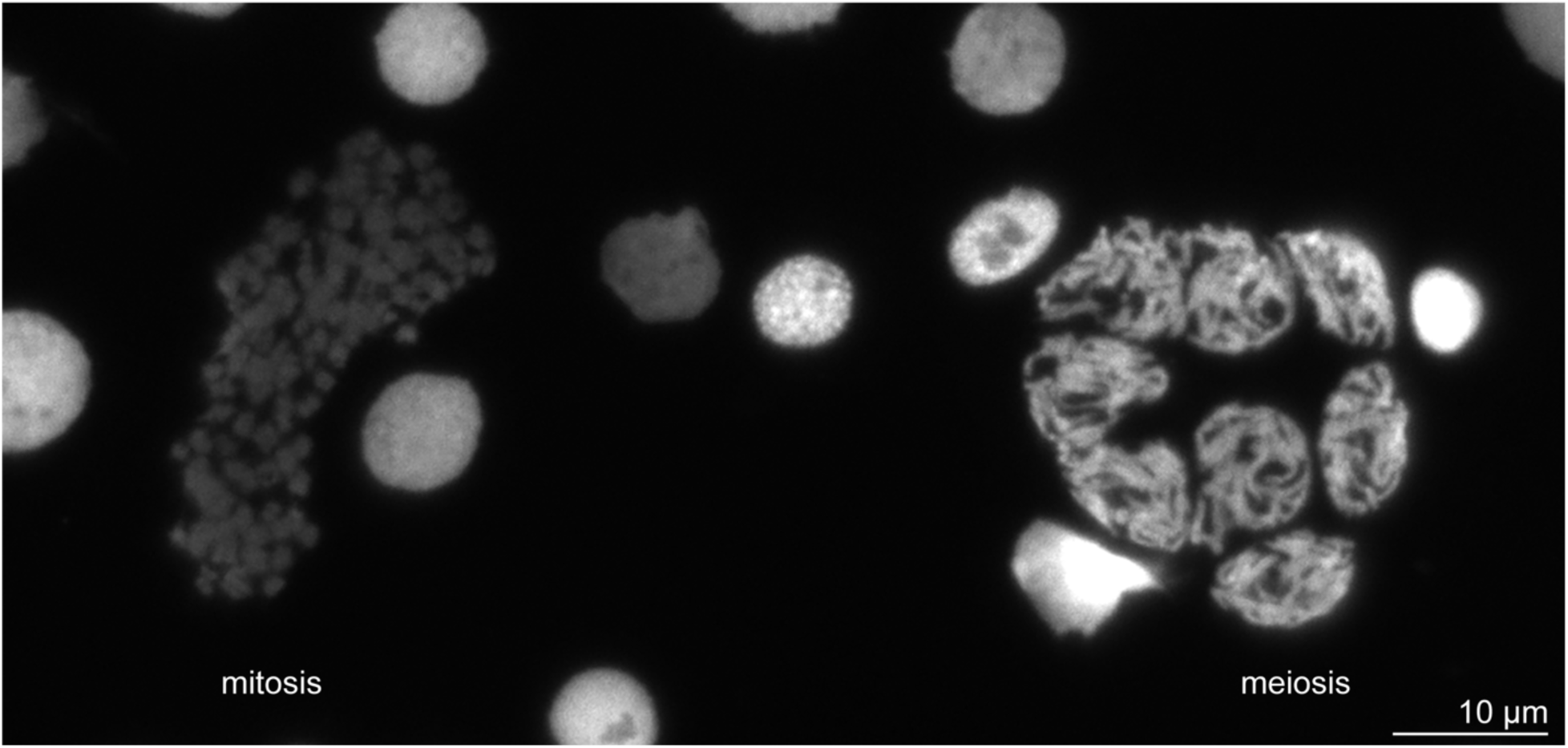
Mitotic and meiotic nuclei in early 5^th^ instar larval testes squash. Representative image showing a cluster of mitotic chromosomes (left) and a cluster of meiotic prophase I cells (right) labeled with DAPI.

**Figure S3.**
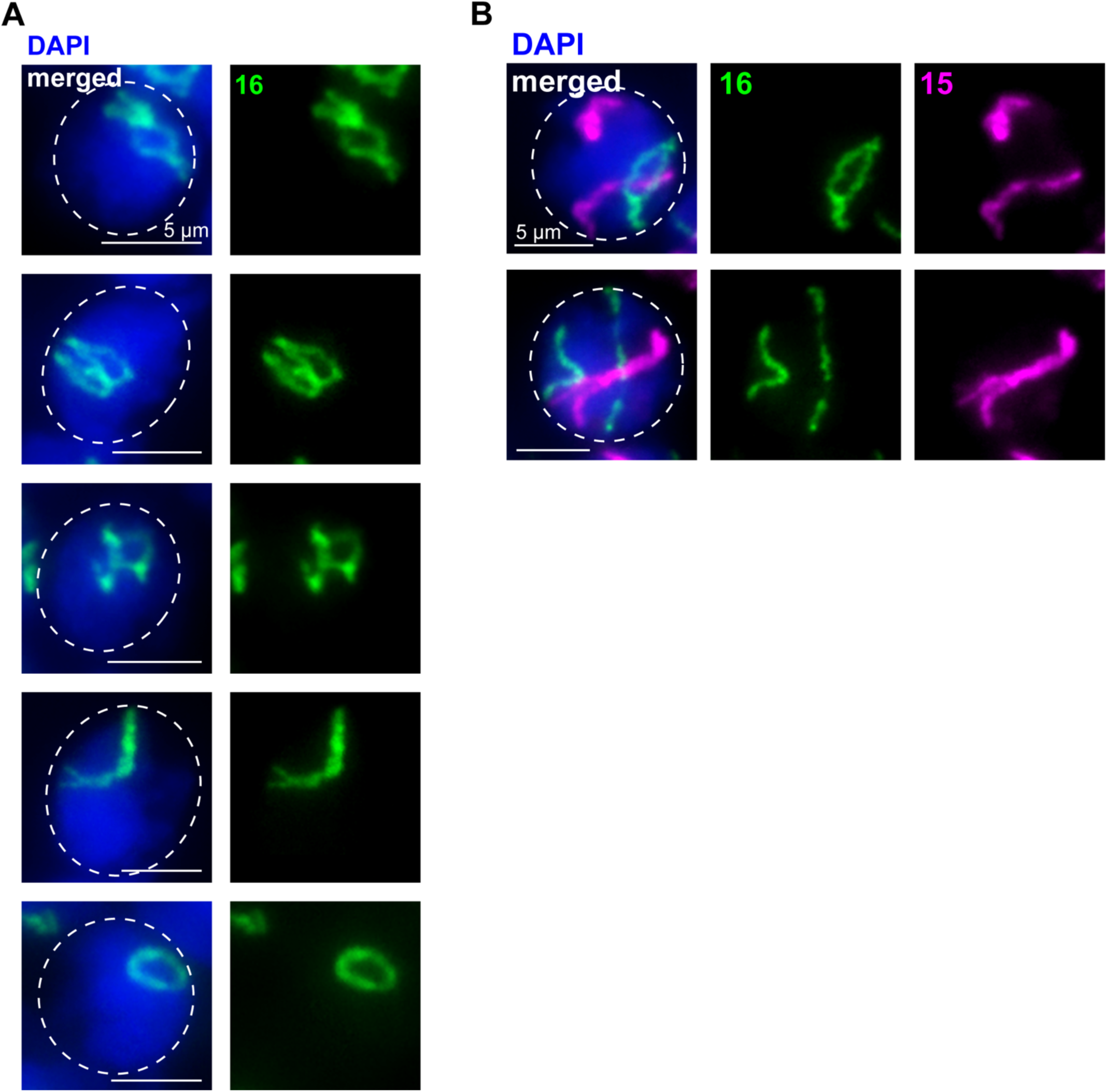
**Partially paired chromosome configurations in early meiotic prophase in larval testes.** A) Oligopaints for chromosome 16 (green) in representative zygotene nuclei. Dashed line approximates the nuclear edge. B) Oligopaints for chromosomes 16 (green) and 15 (magenta) in representative zygotene nuclei. Top: Ch16 has begun pairing while ch15 remains entirely unpaired. Bottom: Ch15 is nearly completely paired while ch16 remains unpaired. Dashed line approximates the nuclear edge.

**Figure S4.**
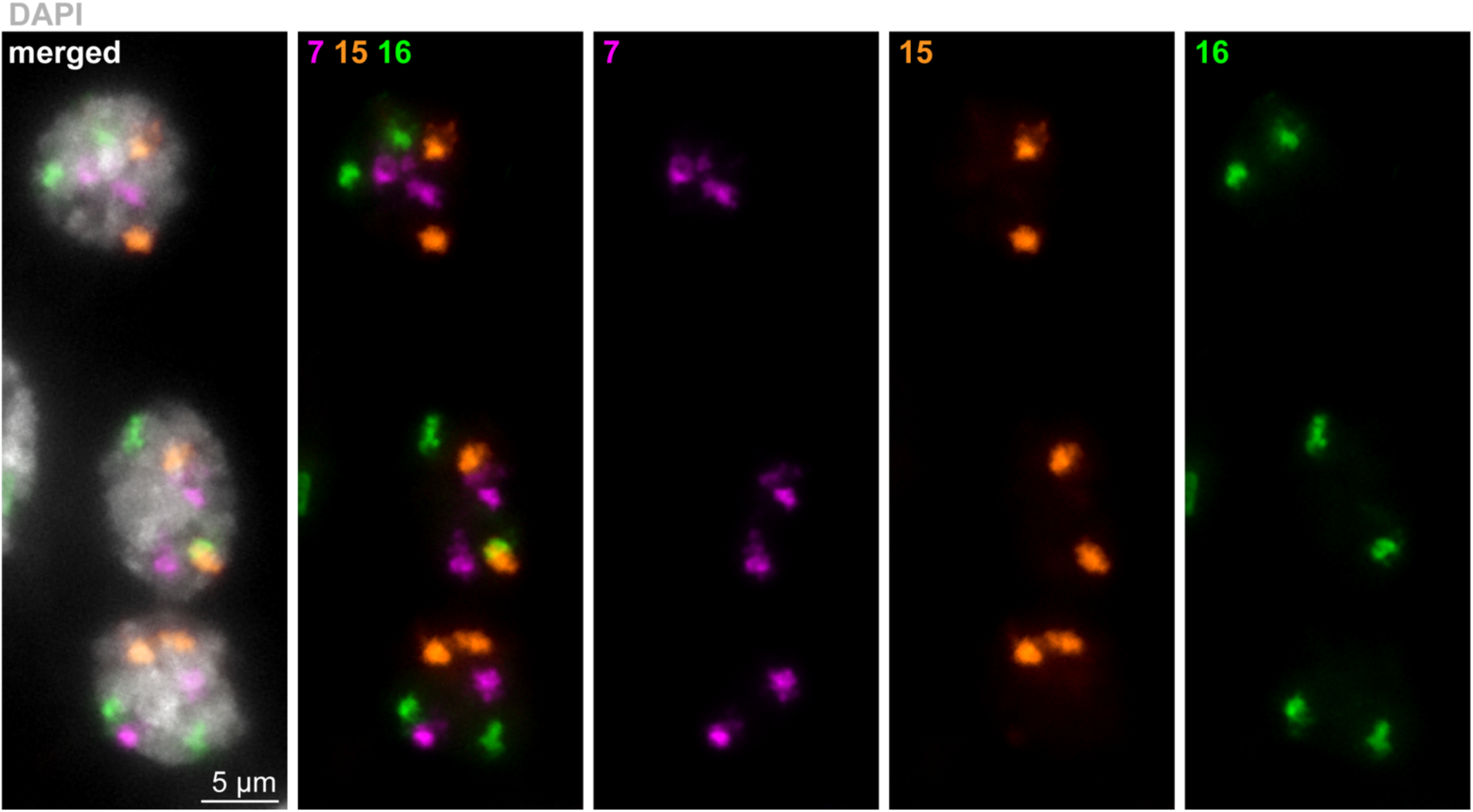
Somatic cells with unpaired homologs from larval testes squashes. Three nuclei (DAPI in gray) labeled with Oligopaints to chromosomes 7 (magenta), 15 (orange), and 16 (green) in somatic cells from late 4^th^ instar larval testes squash. Two signals per nucleus indicates unpaired homologs.

**Figure S5.**
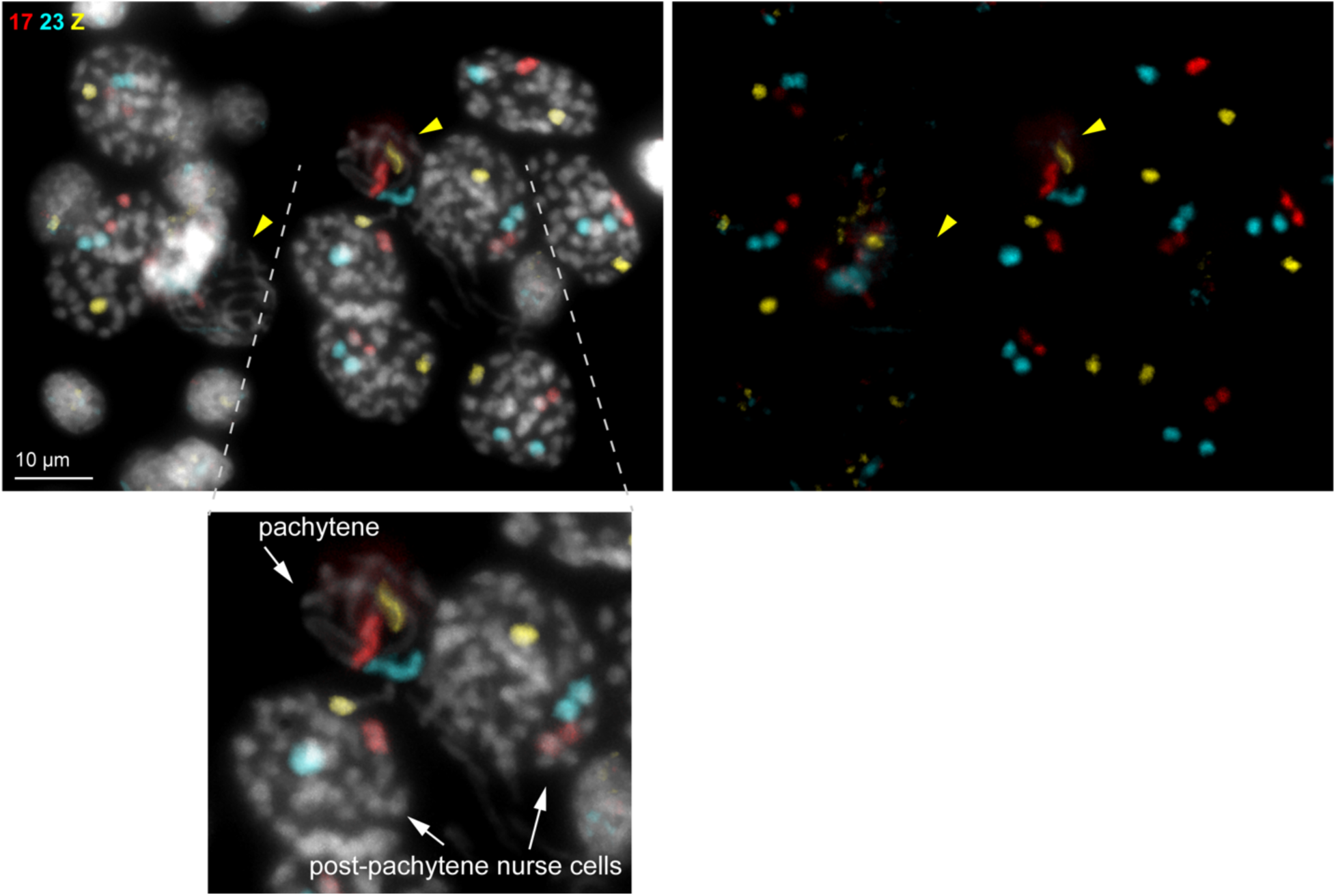
Pachytene and post-pachytene nurse cells labeled with whole chromosome paints. Top: Representative field from early 5^th^ instar larval ovary squash labeled with whole chromosome paints for ch17 (red), ch23 (cyan), and chZ (yellow). Left, merged with DAPI, right, paints alone. Yellow arrowheads indicate pachytene cells. Note: female *B. mori* are heterogametic and harbor a single Z and a single W chromosome (compared to two copies of ch17 and ch23). As the W chromosome is largely repetitive, it is not suitable for the Oligopaint design utilized here. Bottom: zoom showing pachytene and post-pachytene nurse cells, as indicated.

**Figure S6.**
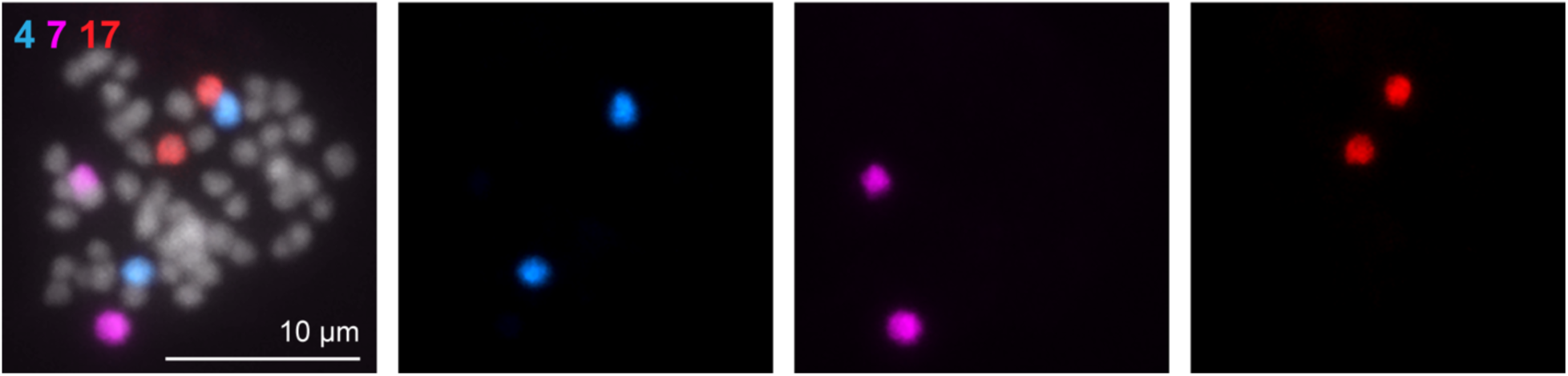
Mitotic cell labeled with whole chromosome paints from 5^th^ instar larval testes. Representative mitotic cell labeled with whole chromosome paints for ch4 (blue), ch7 (magenta) and ch17 (red).

**Figure S7.**
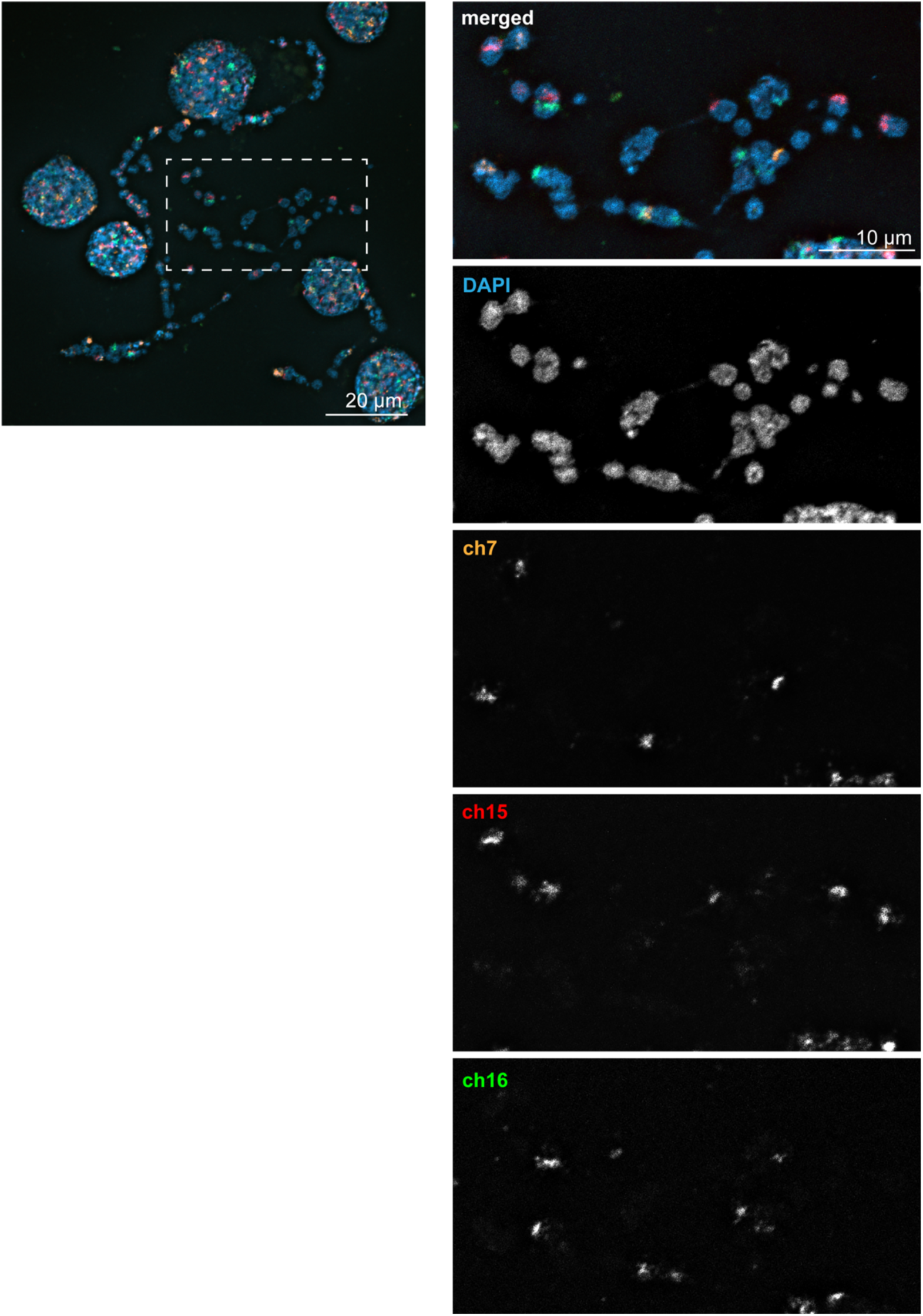
Mitotic spreads from BmN4 cultured cells labeled with whole chromosome Oligopaints. Left: representative mitotic chromosome spread from BmN4 cultured cells labeled with whole chromosome paints for ch7 (orange), ch15 (red), and ch16 (green). White box indicates zoom shown to the right. No entire chromosomes are labeled in the cell line and instead several chromosomes are partially labeled with each paint, indicating a large amount of translocations in the this cell line compared to animals.

**Figure S8.**
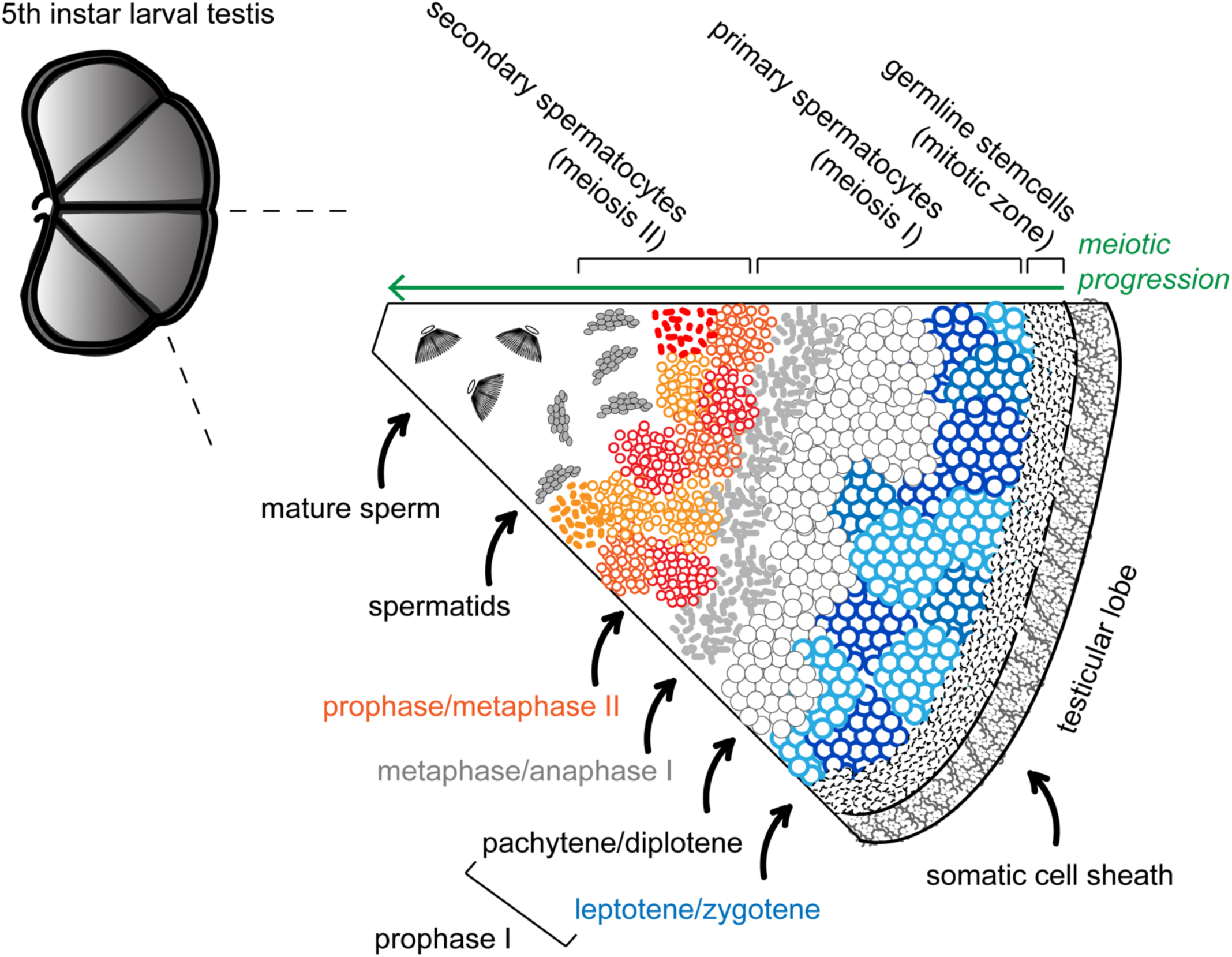
Cartoon schematic of 5^th^ instar larval testis. Mature 5^th^ instar larval testes are comprised of four testicular lobes, each of which harbors germline stem cells (mitotic zone) and spermatocytes in all stages of meiosis up to mature sperm; progressing from right to left in the image. Additionally, each lobe is surrounded by somatic cells in the sheath and in the septae separating the lobes.

**Figure S9.**
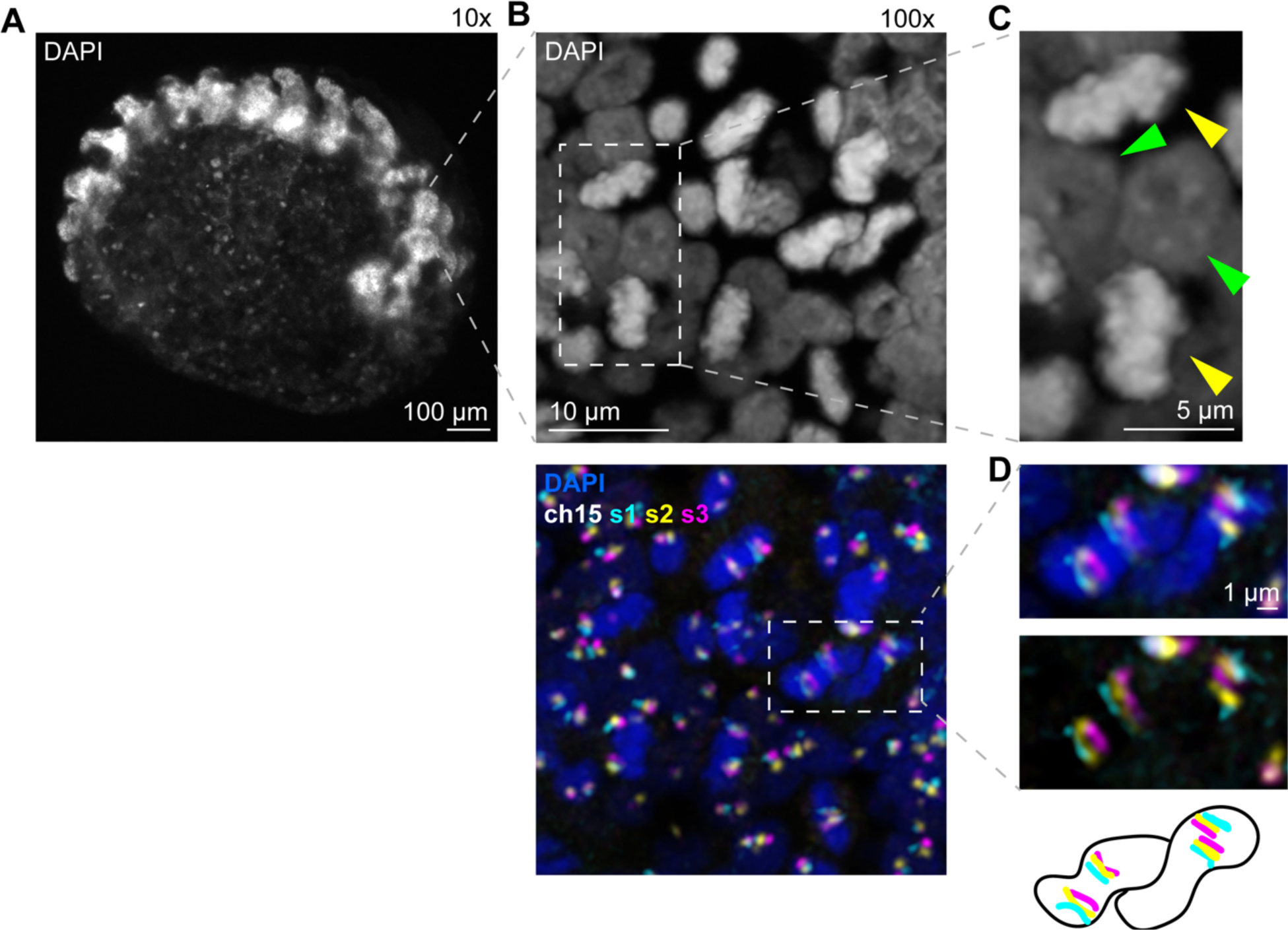
**Ch15 stripe paints in a post-diapause embryo.** A) Whole-mount embryo stained with DAPI imaged at 10x. B) 100x image of whole-mount embryo stained with DAPI (top) and labeled with ch15 stripe paints (bottom). C) Zoom of B (top). Yellow arrow heads indicate mitotic cells, green arrow heads indicate interphase cells. D) Zoom of B (bottom), showing two mitotic cells labeled with ch15 stripe paints and DAPI stain (top) or only stripe paints (middle). Bottom, cartoon schematic, with the black outline representing the border of the DAPI stain.

**Figure S10.**
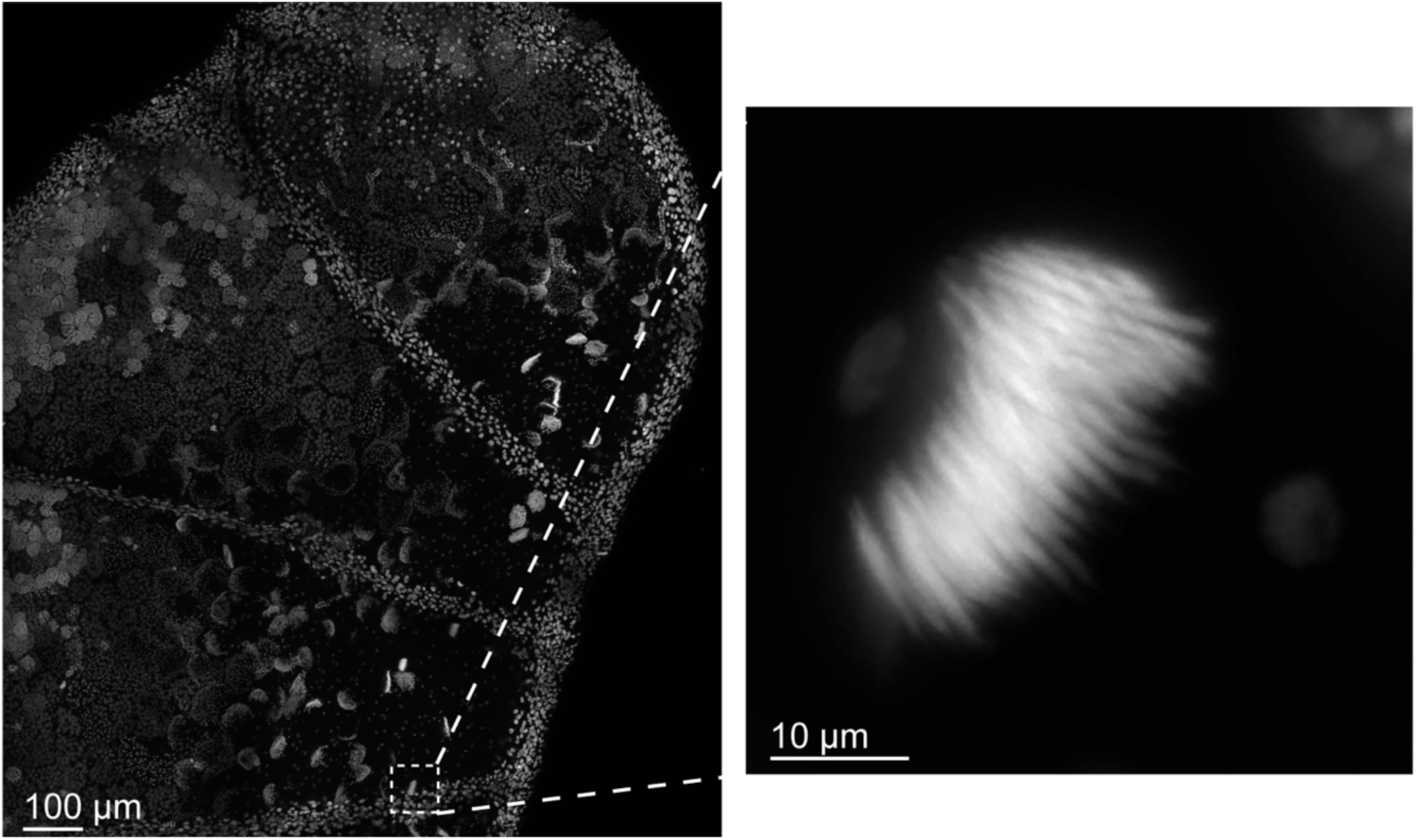
Mature sperm in 5^th^ instar larval testes. Left: 10x image of larval testes stained with DAPI. White box indicates zoom shown to the right. Right: zoom of mature eupyrene sperm bundle labeled with DAPI.

**Figure S11.**
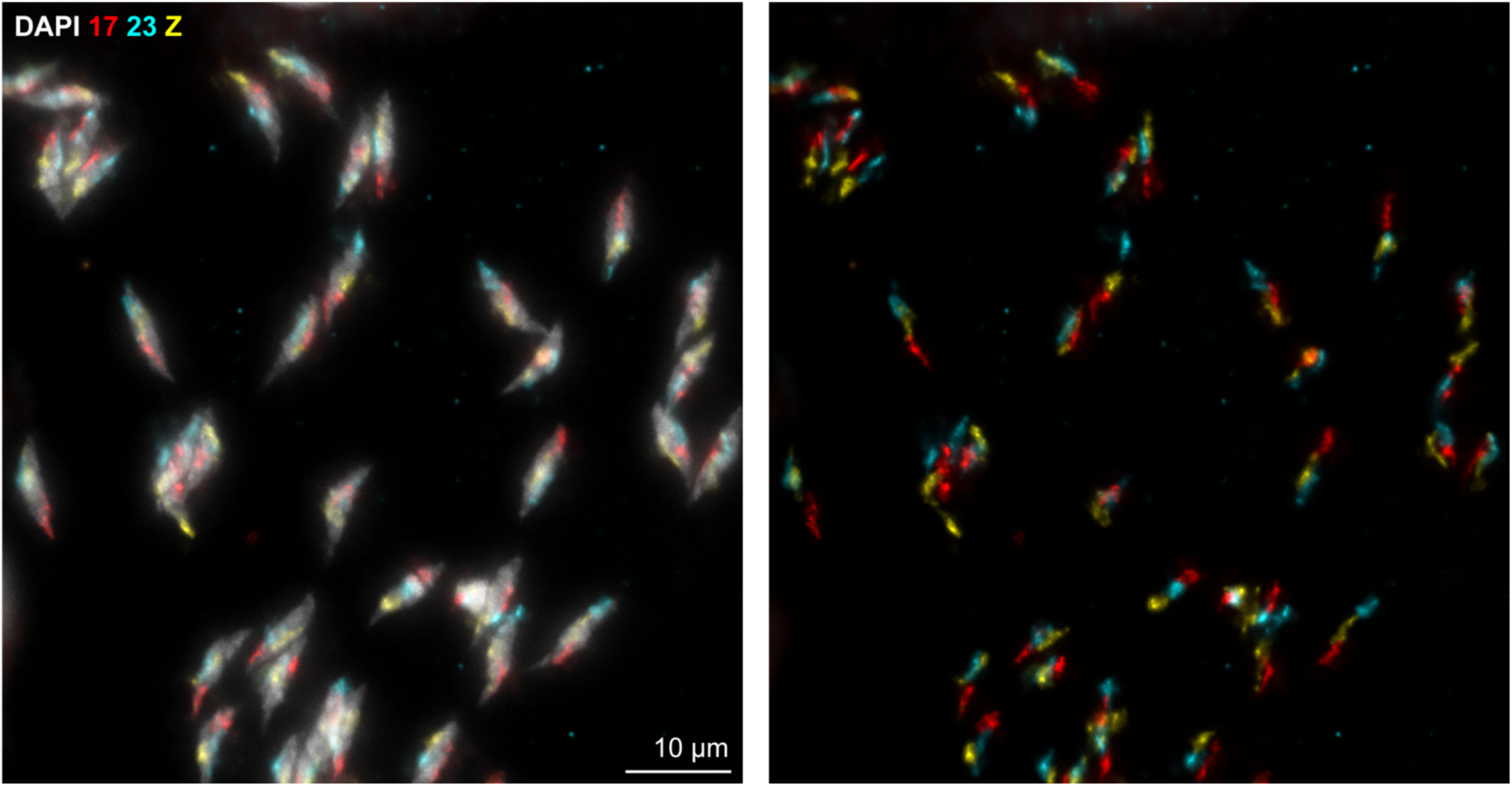
Oligopaints in mature sperm from 5^th^ instar larval testis squash. Oligopaints labeling chromosomes 17 (red), 23 (cyan) and Z (yellow) in mature sperm from a 5^th^ instar larval testis squash. DAPI is shown in gray.

**Figure S12.**
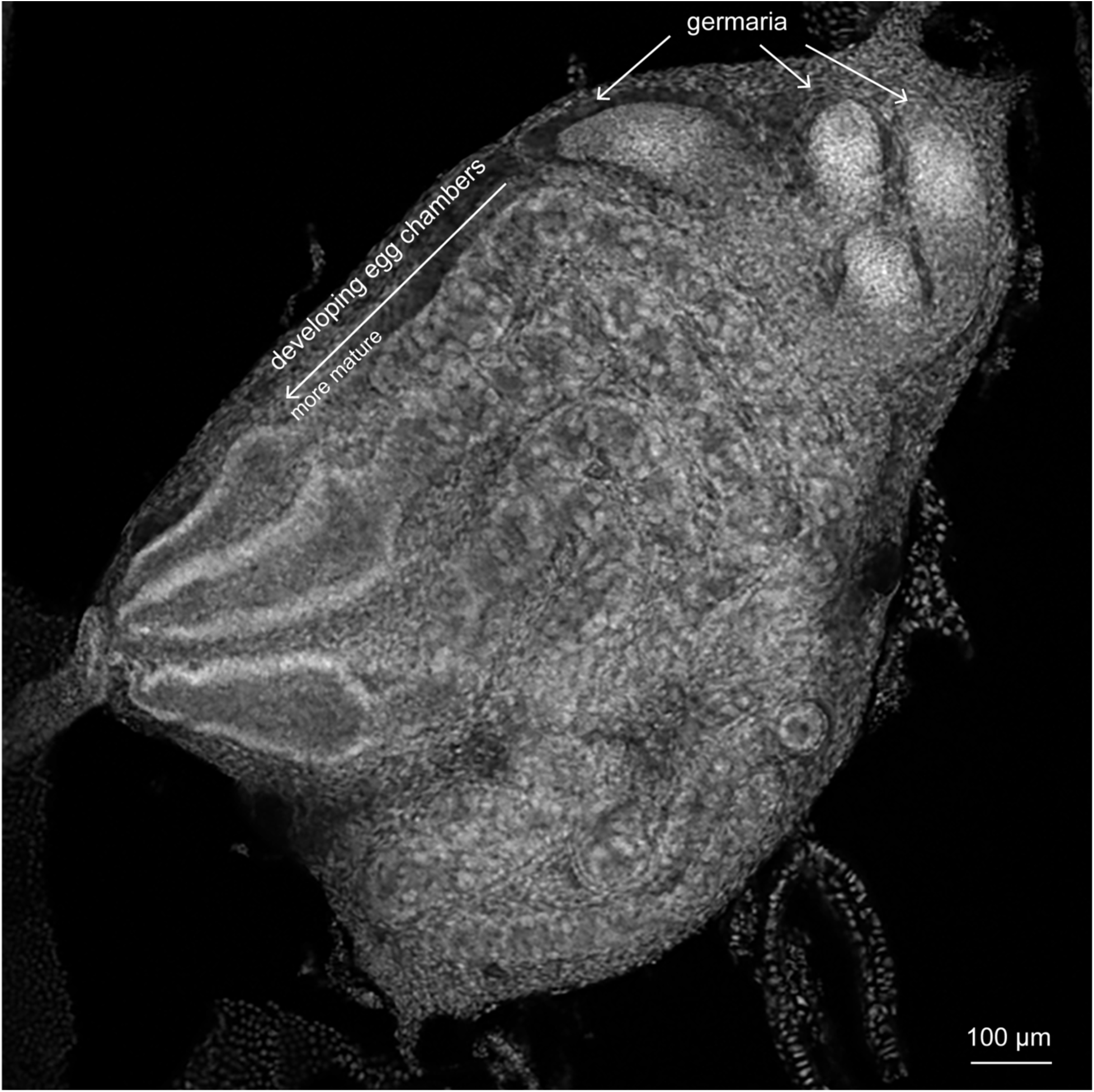
5^th^ instar larval ovary. DAPI staining on a whole mount 5^th^ instar larval ovary.

## Notes

### Competing Interest Statement

The authors have declared no competing interest.

